# Differential co-localisation analysis of multi-sample and multi-condition experiments with spatialFDA

**DOI:** 10.64898/2026.04.13.718197

**Authors:** Martin Emons, Fabian Scheipl, Samuel Gunz, Elizabeth Purdom, Mark D. Robinson

## Abstract

Advances in spatial omics data generation have led to an explosion in new datasets that record the spatial location of transcripts and proteins. However, challenges remain in the analysis of spatial omics data. One important analysis is differential cellular co-localisation (CCoL): the quantification of the clustering, or spacing, of one or more cell types across multiple conditions. Our framework `spatialFDA` combines methodology from spatial statistics with functional data analysis to accurately quantify and test for differences between conditions in CCoL across spatial scales. Using two simulation studies, we show that `spatialFDA` performs well in controlled settings. Furthermore, `spatialFDA` recovers known biological processes in type-1 diabetes and adds insights about the CCoL strength in space. `spatialFDA` is readily available as an open-source `Bioconductor R` package.

## 1 Introduction

Recent advances in the field of spatial omics allow for the high-dimensional molecular characterisation of cells in their native biological context [1, 2]. Cells can be profiled spatially via spatial proteomics (IMC, MIBI-TOF, 4i, CODEX [3, 4, 5, 6]), spatial transcriptomics (10x Visium, Visium HD, CosMx, Xenium [7, 8, 9, 10]) and spatial epigenomics [11]. One biological question that can be elucidated with spatial omics technologies is cellular co-localisation (CCoL), which refers to finding cell types that are more or less often in the vicinity of other cell types than by chance. This definition of cellular co-localisation is closely related to the spatial statistics concepts of clustering and spacing. Clustering is a form of positive co-localisation in comparison to a random pattern, whereas spacing corresponds to negative co-localisation. A related concept to CCoL from bioinformatics is called spatial neighbourhood analysis, which defines a fixed neighbourhood and quantifies, for example, changes in cell type proportions in neighbourhoods across conditions [12, 13, 14]. Neighbourhood analysis can also be performed across varying neighbourhood sizes, reducing the gap to classical spatial statistics approaches [15]. CCoL can be observed in the tumour micro-environment, where tumour-associated macrophages can either promote or inhibit tumour growth [16]. Furthermore, the degree of CCoL can differ between conditions, such as that observed across cold, hot and mixed tumour subtypes with implications for clinical outcomes [4]. We call this analysis of CCoL across conditions differential CCoL.

One way to quantify CCoL is with point pattern analysis [17], which approximates spatial omics data as points. First, the cellular boundaries are segmented and all features within a cell are recorded in order to give the cell a categorical identity, for example, a cell type. Achieving accurate cell segmentation is a challenging problem itself, for example, mapping measurements of transcripts in three dimensions into two [18, 19, 20]. The cell is then approximated as a point via the cell centroid, and the cell type is added as a *mark*, i.e. a categorical or continuous feature recorded per cell.

The degree of CCoL of points with a cell type mark (or across pairs of cell types) is quantified with point pattern metrics for correlation and spacing. These quantifications are often spatial metrics such as Ripley’s *K* function, Besag’s *L* function or the nearest-neighbour function *G* [17, 21, 22, 23]. These spatial metrics are based on the notion of quantifying a measure in an *r*-neighbourhood around a point of interest. By expanding the radius *r*, the metrics can be represented as a function of the radius. Repeating this process for every cell in the query cell type, spatial metrics represent estimates averaged over an entire field of view (FOV) for a given cell type. The estimation of the FOV can itself be challenging in the presence of tissue artifacts such as folds or missing regions. These apparent technical inhomogeneities can lead to biased spatial metrics estimations [24, 25, 26]. Typically, the spatial metrics are then normalised by the number of points per FOV, *n*, and the size of the FOV, |*W* |, making them comparable across FOVs of differing sizes and numbers of points [24], [17, p. 204].

Calculating one spatial metric per FOV raises the question of how to compare them in experiments with multiple FOVs, samples and conditions. These different levels of variability should be taken into account in differential CCoL analysis. Failure to do so can result in overly optimistic results. In the case of a scalar response, datasets with these differing levels of variability can be analysed well with linear mixed-effects models [27, 28]. The challenge in spatial data becomes how to optimally summarise the functional spatial metrics in a single number. One option is to calculate the area-between-curves (ABC) of a spatial metric (e.g., to a completely spatially random baseline) and compare the ABC between conditions while accounting for non-independent FOVs. A variant of ABC (see Methods 5.5) is at the core of two competing differential CCoL packages `spicyR` and `smoppix` [29, 30].

We hypothesise that the compression of a function into a single number may come at a loss of sensitivity, based on the fact that two rather disparate functions can have the same ABC. Meanwhile, the field of functional data analysis (FDA) now provides many flexible tools to handle functional responses and covariates [31]. Thus, instead of condensing spatial metrics into scalars, FDA can be used to compare the entire function over its domain *r*. Two existing methods compare entire spatial metrics across conditions using FDA. The first method, `SpaceANOVA`, models the spatial metric with a one-way or two-way functional ANOVA [32]. The other method, `mxfda`, uses the spatial metric as a covariate to model survival, binary and continuous outcomes, a modelling setup called scalar-on-function regression [33, 34, 35] (more details in Methods 5.6). However, `SpaceANOVA` does not allow nested variability to be modelled explicitly and `mxfda` only offers a setup to add the spatial metric as a functional covariate and not as outcome in the regression model.

In this paper we present `spatialFDA`, a comprehensive `R Bioconductor` package to quantify differential CCoL using function-on-scalar regression with a functional additive mixed-effects model framework, a novel methodological approach that has not yet been explored for spatial omics data analysis [36]. The new statistical inference framework of `spatialFDA` allows users to define differing layers of variability while retaining the scale information of the spatial metrics. We show sensitive and calibrated performance of `spatialFDA` across two simulation studies and recover differential CCoL using a small cohort study of type-1 diabetes as well finding new patterns of differential CCoL in a large non-small cell lung cancer dataset.

## 2 Results

### 2.1 spatialFDA unifies multi-condition comparisons of spatial metrics

`spatialFDA` takes as input the measurements from sub-cellular spatial omics technologies with the aim of detecting *differential CCoL* of cells/features in space. As such, cells are approximated by their cell centroid and are given marks, most often a cell type label (Figure 1A). In complex experimental setups, there can be multiple FOVs per condition and sample (Figure 1B). Spatial metrics measuring CCoL can be calculated for each FOV, and most spatial metrics from point pattern analysis are functions of radius *r*. That is, for a given point in space, (*x, y*)_*i*_, an *r*-neighbourhood surrounding the point is constructed and (spatial) characteristics of this *r*-neighbourhood can be quantified (e.g., average correlation or spacing) [17, p. 204],[24]. Thus, we have multiple spatial metrics that are functions of *r*, each corresponding to a FOV (Figure 1C).

**Figure 1:**
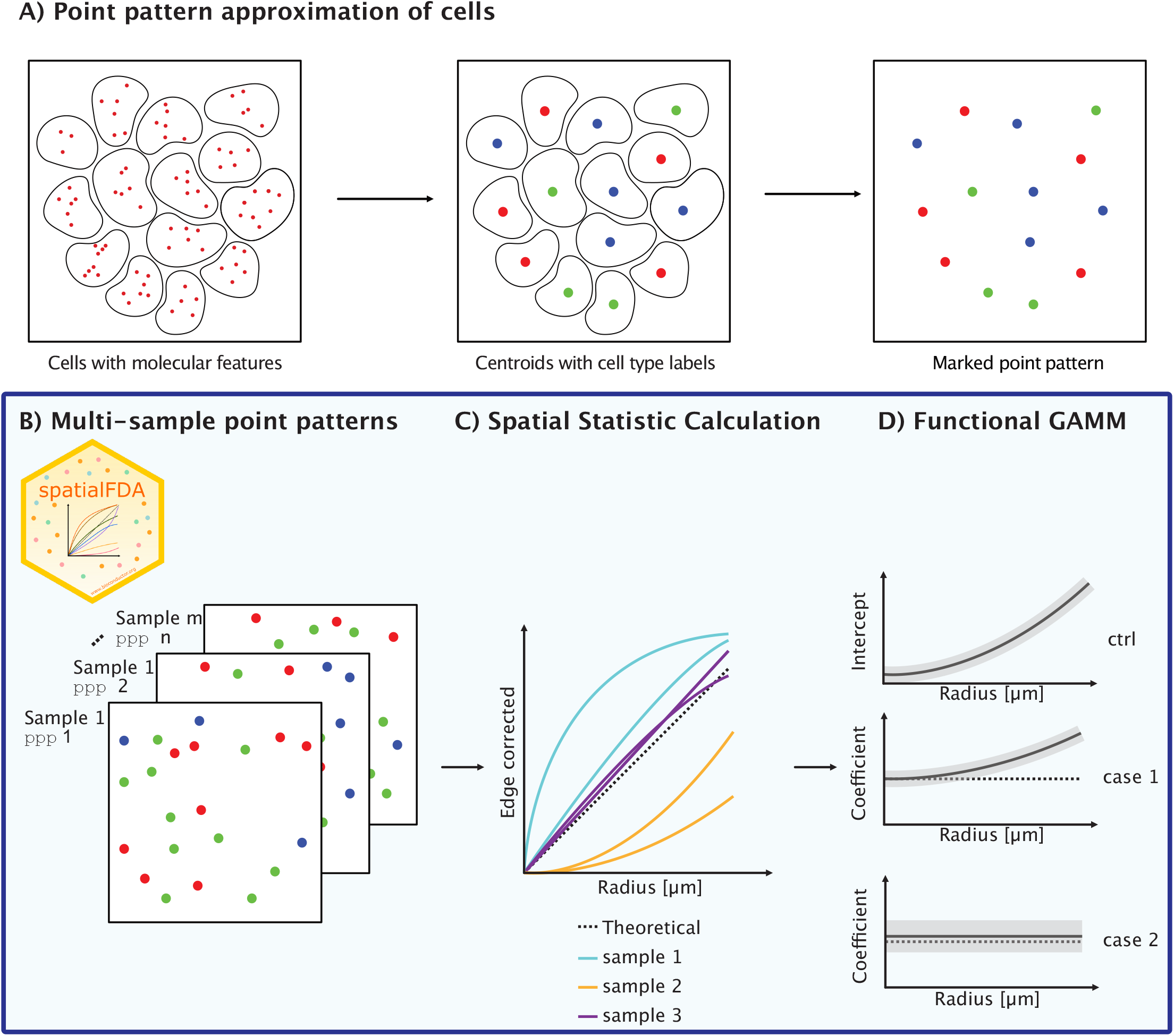
Summary of the functionality of `spatialFDA`. **A** Spatial omics data can be represented as points via, for example, cell centroids. The mark added is the cell type according to the molecular profile of the cell. **B** The data stored as `SpatialExperiment` are converted to a planar point pattern (`ppp`) [44] data representation. In multi-sample experiments, this can lead to several point patterns per sample (e.g., different FOVs) and multiple samples per experimental condition. **C** The `ppp` objects are the input for the spatial metric calculation. These are functions of radius *r* (typically corrected for edge-effects). Most often, these functions are calculated for each FOV. **D** The spatial metrics are further analysed with a fGAMM [45, 46, 36]. The control group functions are estimated as an intercept curve and the case group functions are estimated as difference coefficients. In this example, case 1 shows an increased CCoL at larger radii, whereas case 2 shows no difference in CCoL across the domain *r*.

In order to compare the spatial metrics between conditions, `spatialFDA` uses *functional* generalised additive mixed models (fGAMM), which accommodate flexible experimental designs (i.e., covariance structures). `spatialFDA`’s fGAMM performs inference on the entire spatial metric to compare groups of functions (e.g., from case vs. control samples) while accounting for random effects between samples and FOVs (see Methods 5.1.2). This generates model coefficients that are interpretable across spatial scales (see Figure 1D) [37, 36, 38]. In order to get a well-calibrated estimator of differential co-localisation, `spatialFDA` corrects post hoc for model residuals that are neither independent nor identically distributed via cluster-corrected covariance matrices, an approach novel to differential co-localisation analysis [39, 40, 41]. The application of post hoc cluster-correction to FDA makes these methods more generalisable to other constrained functional data types than classical FDA approaches are intended for.

In addition to the readout of the functional model coefficients, the fGAMM provides an omnibus *F*-test result over the entire domain. This test gives an indication of *any* CCoL difference between case and control spatial metrics along the domain *r*. Individual pointwise confidence bands are also provided at each *r*, but can only be used as an exploratory result and not for statistical inference at a specific radius *r*.

`spatialFDA` is provided as a `Bioconductor R` package, allowing a simple integration into existing spatial analysis workflows [42, 43].

### 2.2 Simulation study shows a well-calibrated performance of spatialFDA

We performed a simulation study to investigate the ability of `spatialFDA` to detect changes in CCoL. The simulation design was adapted from Baker et al. (2023) and serves as a way to understand performance for both true null experiments and controlled changes [47]. The general simulation setup is described in detail in Section 5.2 (see also Figure S1). First, empty tissue scaffolds are created by packing random-radii (*r*) circles to mimic the density of a tissue slice (Figure S1A). From these circles (cells), a graph is constructed with centroids as nodes and edges between touching circles. Using this graph, the centroids of the simulated circles are annotated with one of four cell type marks (Figure S1B,C). For *K* cell types, the cell type assignment takes as input a vector with the specified cell type proportions *p* ∈ ℝ^*K*^ and a matrix *H* ∈ ℝ^*K*×*K*^ defining the interaction probabilities between the cell types (see Methods 5.2).

In any simulation scenario, we only modulated the interaction probabilities of cell type 0 with itself (i.e., *p*_0_ is fixed and *H*_0,0_ differs across conditions). Specifically, *H*_0,0_ was sampled from a truncated [0,1] normal distribution with a condition-specific mean and fixed standard deviation. To generate additional variation, we added a sample-specific offset to the condition-wise mean (see Methods 5.2). From the larger simulated tissue section, we selected smaller FOVs (see Figure S1C). For each condition, we defined 5 samples, sampling 10 images per sample (resulting in a total of 50 images per condition). Example simulated point patterns are shown in Figure S2 and calculated spatial metrics for one realisation of the simulation can be found in Figure S3. The entire simulation (Figure S1) was repeated 500 times to obtain an estimate of the power and calibration of the methods to discriminate between true positives and true negatives. Despite a parametrisation that fixes the cell type proportion vector *p*_0_ across conditions and only changes the interaction probability *H*_0,0_, the observed average cell type intensities (normalised proportions) are drastically changed (Figure S4). This suggests that the simulation framework does not clearly distinguish between differences in CCoL and proportions.

Further simulation scenarios were generated by changing the (constant) cell type proportions *p*_0_ (the different panels in Figure 2). Each scenario allowed us to compare the performance of `spatialFDA` to the four competitor methods `spicyR`, `smoppix`, `SpaceANOVA` and `mxfda` plus a baseline across different settings [29, 30, 32, 33]. As a baseline, we compared the average *intensities* (number of cells of type 0 per image divided by the image area |*W* |, *λ* = *n*(*x*_0_)*/*|*W* |). This is a non-spatial metric, comparing the average cell type proportion across images. The intensities were aggregated into a vector across samples and compared between conditions with a linear mixed model (see Methods section 5.5 and 5.6).

**Figure 2:**
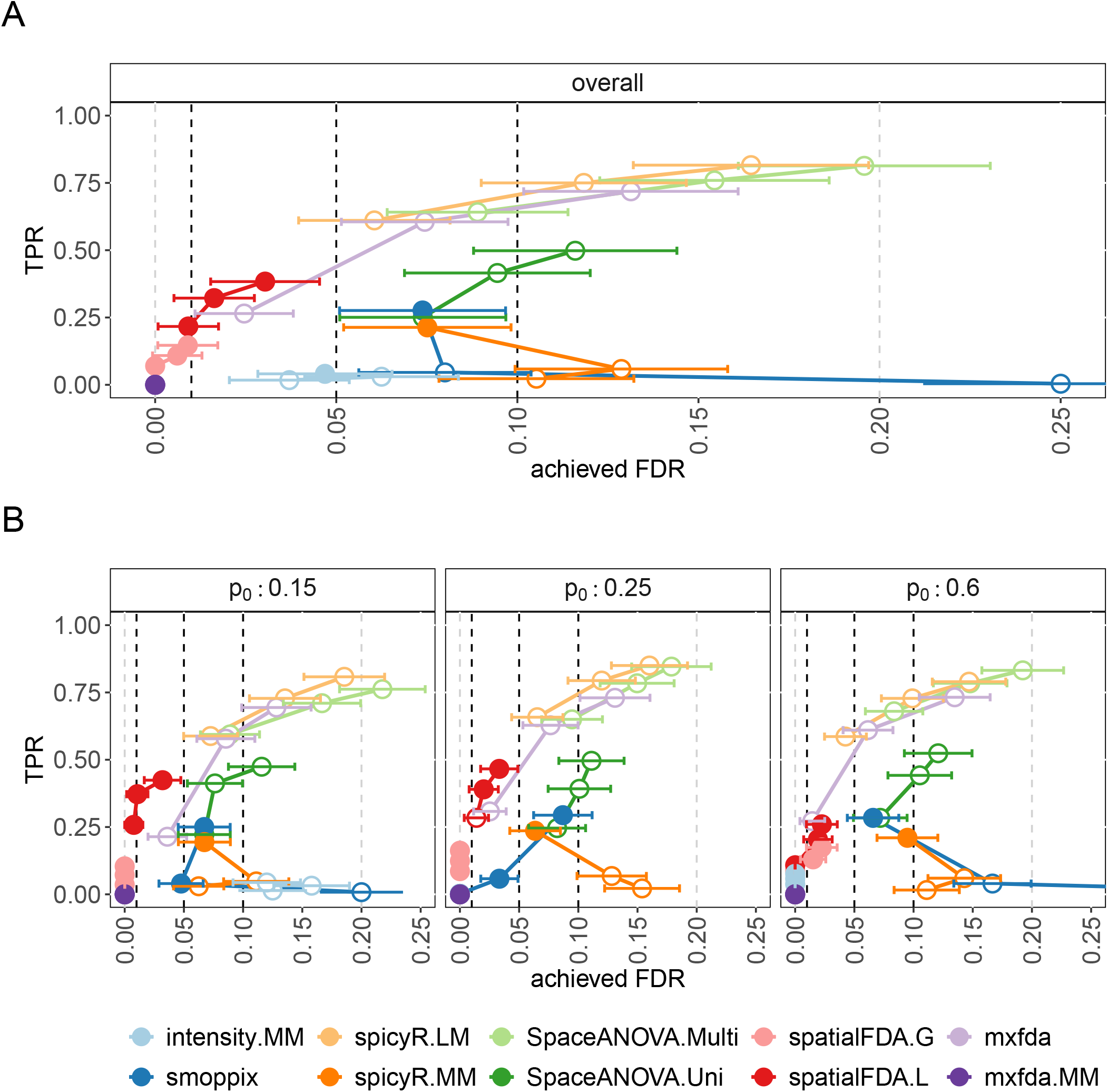
Assessment of the calibration of the methods across different proportions. The curves plot the true-positive rate (TPR) against the achieved false discovery rate (FDR). The black dashed lines indicate the nominal/target FDR at conventional cut-offs (FDR = 0.01, 0.05 and 0.1). The three points per method correspond to the observed FDR for the three targeted nominal FDR cut-offs. If the achieved FDR is smaller than the nominal FDR cut-off, i.e., the target FDR is met by the statistical test, the points are filled; otherwise they are open. The error bars indicate Wald confidence interval of the Monte Carlo standard errors (MCSE) per observed FDR. **A** Overall performance results across all simulations. **B** Performance results split by the prior cell type proportion of the perturbed cell type.

The simulation study results for the various flavours (Table 1) are summarised in Figure 2; to assess the trade-off between sensitivity and error control, we focused on true-positive rate (TPR) vs. achieved false discovery rate (FDR). Curves were generated by mixing null settings (i.e., in expectation 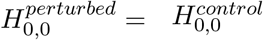) with those from induced differences (in expectation 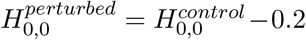) in CCoL at equal proportions (see Methods 5.2). Since the FDR can be sensitive to the exact proportion of true negatives and true positives, we also present receiver operating characteristic (ROC) curves (Figure S6).

**Table 1:** Methods included in the simulation study. The baseline is a linear mixed model comparing the average intensity (number of cells divided by the area of the image) of a cell type between images. The scalar methods include `spicyR` and `smoppix`. Existing functional methods are `SpaceANOVA` and `mxfda` as well as our proposed method `spatialFDA`. For all methods we test different variants and testing options.

| Method | Quantification | Comparison |
| --- | --- | --- |
| `intensity.MM` | average intensity | linear mixed model |
| `spicyR.LM` | ABC of $L$ function | linear model |
| `spicyR.MM` | ABC of $L$ function | linear mixed model |
| `smoppix` | ABC of $G$ function | linear mixed model |
| `SpaceANOVA.Uni` | Derivative of $K$ function | one-way functional ANOVA |
| `SpaceANOVA.Multi` | Derivative of $K$ function | two-way functional ANOVA |
| `mxfda` | $K$ function | functional logistic regression |
| `mxfda.MM` | $K$ function | functional logistic mixed-effects regression |
| `spatialFDA.L` | $L$ function | functional mixed model |
| `spatialFDA.G` | $G$ function | functional mixed model |

Our proposed method `spatialFDA` shows the most consistent conservative calibration while still retaining an acceptable TPR. `spatialFDA.L` is well calibrated across all simulation scenarios with the highest TPR relative to FDR, whereas `spatialFDA.G` shows a lower TPR on average while also maintaining good calibration. We note that our novel use of cluster-corrected covraince is critical in providing effective calibration of *p*-values when applying traditional FDA strategies to spatial metrics (Figures S7 and S8).

In general, Figure 2A highlights the necessity of mixed-model frameworks for complex spatial omics datasets. All methods that do not account for nested covariance structures are severely miscalibrated (`spicyR.LM`, `SpaceANOVA.Uni, mxfda`).

The scalar differential CCoL methods which account for repeated measurements, `spicyR.MM` and `smoppix`, control type I errors better for the nominal 10% FDR cut-off but at the cost of sensitivity. In the high proportion simulations (panel *p*_0_ = 0.6 in Figure 2B) both `spicyR.MM` and `smoppix` show strongly inflated type I errors.

Both variants of the functional method `SpaceANOVA` are not well calibrated, showing strong increases in FDR. Interestingly, this increase in FDR is stronger for the two-way ANOVA model `SpaceANOVA.Multi` than for the one-way ANOVA model `SpaceANOVA.Uni`. An adjustment to a logistic regression with mixed-effects (`mxfda.MM`) was not successful and led to no detections, most likely due to an implementation problem.

The non-spatial baseline (`intensity.MM`) is generally not well calibrated in comparison to the nominal FDR cut-offs (Figure 2A). This points to a general conceptual question of whether differences detected are due to *intensity* (number of cells normalised by the area) or CCoL (re-arrangement of cells without changing the intensities of cells).

Taken together, `spatialFDA` is a conservative estimator of CCoL but shows decent TPR in comparison to the competitor methods. Furthermore, accounting for repeated measurements with mixed-effects is necessary in complex spatial omics datasets. The scalar mixed-effects methods (`spicyR.MM` and `smoppix`) show reduced statistical power. The functional methods, `SpaceANOVA`, `mxfda`, show inflated FDR in our simulation study rendering conclusions about TPR difficult. In general, cell type intensity is an important baseline to distinguish between changes in *proportions* and *CCoL*. Interestingly, *K* and *L* functions explicitly normalise for differences in non-spatial intensity, whereas the *G* function does not. This means that analyses with *K* and *L* functions will suffer less from the confounding with average intensity whereas *G* functions assess both differences jointly. The choice for users thus depends on whether they regard overall changes in cell type proportions as a spatial effect or as a confounder they want to account for. This guides the use of either *G* (not intensity normalised) or *L* (intensity normalised) functions. In addition, we have added the option to adjust for the intensity covariate in the `spatialFDA.G` fGAMM, trying to disentangle intensity from CCoL differences with *G* functions (Figure S7 and Figure S8).

### 2.3 Simulating CCoL at different scales shows good performance of spatialFDA.L **and** mxfda

In the previous simulation we tried to ensure that parameters like cell spacing in a biological setting were mimicked, but the simulation did not allow us to introduce specific types of changes. Specifically, a motivation for `spatialFDA` was to detect CCoL across a range of radii, but this could not be directly manipulated in the previous simulation. To query this more directly, we ported the simulation developed by Canete et al. (2022). Although less biologically inspired than our simulation, the Canete et al. (2022) simulation has a mechanism to introduce changes in CCoL at specific scales [29]. They simulated as follows (see Supplementary Figure 4 of [29] for details): i) Cells of type *A* are sampled from a random Poisson process with a specified intensity; a density around type *A* cells is estimated using a disc kernel; ii) The locations of cell type *B* are simulated according to a Poisson process with an intensity proportional to the type *A* density. By varying the radius of the disc kernel, the scale of the *A*-*B* CCoL interaction changes. Figure 3 shows TPR vs. achieved FDR curves for the baseline model `intensity.MM`, `spicyR` variants, `smoppix`, `spatialFDA` variants, `SpaceANOVA` variants and `mxfda` variants. All methods were set to evaluate effects across a radius range of *r* ∈ [0, 100]. The corresponding ROC curves are shown in Figure S9.

**Figure 3:**
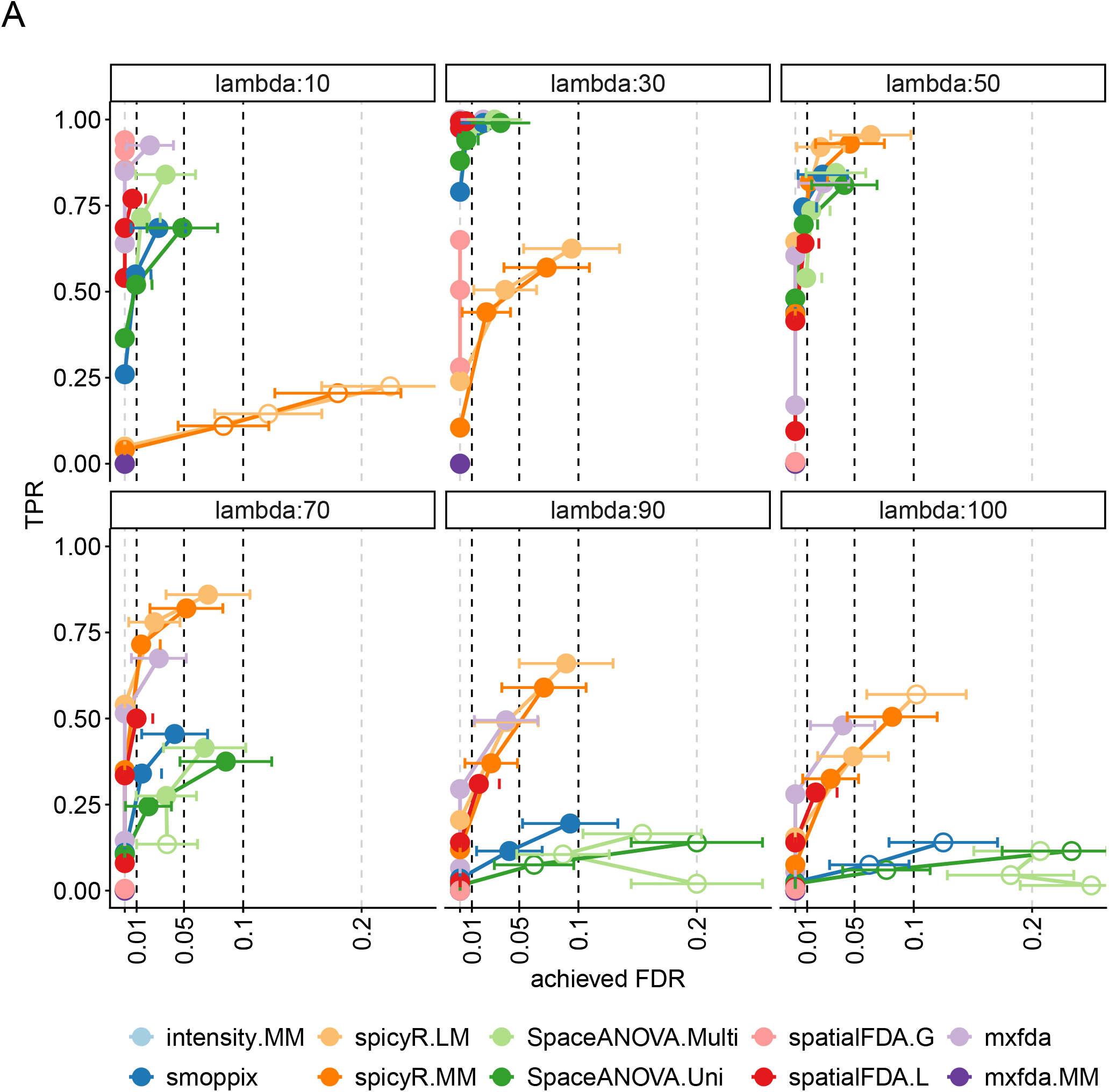
True-positive rate (TPR) vs. achieved false discovery rate (FDR) curves for the simulation from Canete et al. All methods are parameterised to calculate spatial metrics (*L* and *G* curves) on a [0,100] interval and compared by ABC (`spicyR`, `smoppix`) or functional data analysis (`spatialFDA`, `SpaceANOVA`, `mxfda`) as well as a non-spatial baseline (`intensity.MM`). The true length scale of the differential CCoL effect changes between the facets of the plot from 10*µm* to 100*µm* simulated differences in space between the two simulated cell types. The black dashed lines indicate the nominal/target FDR at conventional cut-offs (FDR = 0.01, 0.05 and 0.1). The three points per method correspond to the observed FDR for the three targeted nominal FDR cut-offs. If the achieved FDR is smaller than the nominal FDR cut-off, i.e., the target FDR is met by the statistical test, the points are filled; otherwise they are open. The error bars indicate Wald confidence interval of the Monte Carlo standard errors (MCSE) per observed FDR.

**Figure 4:**
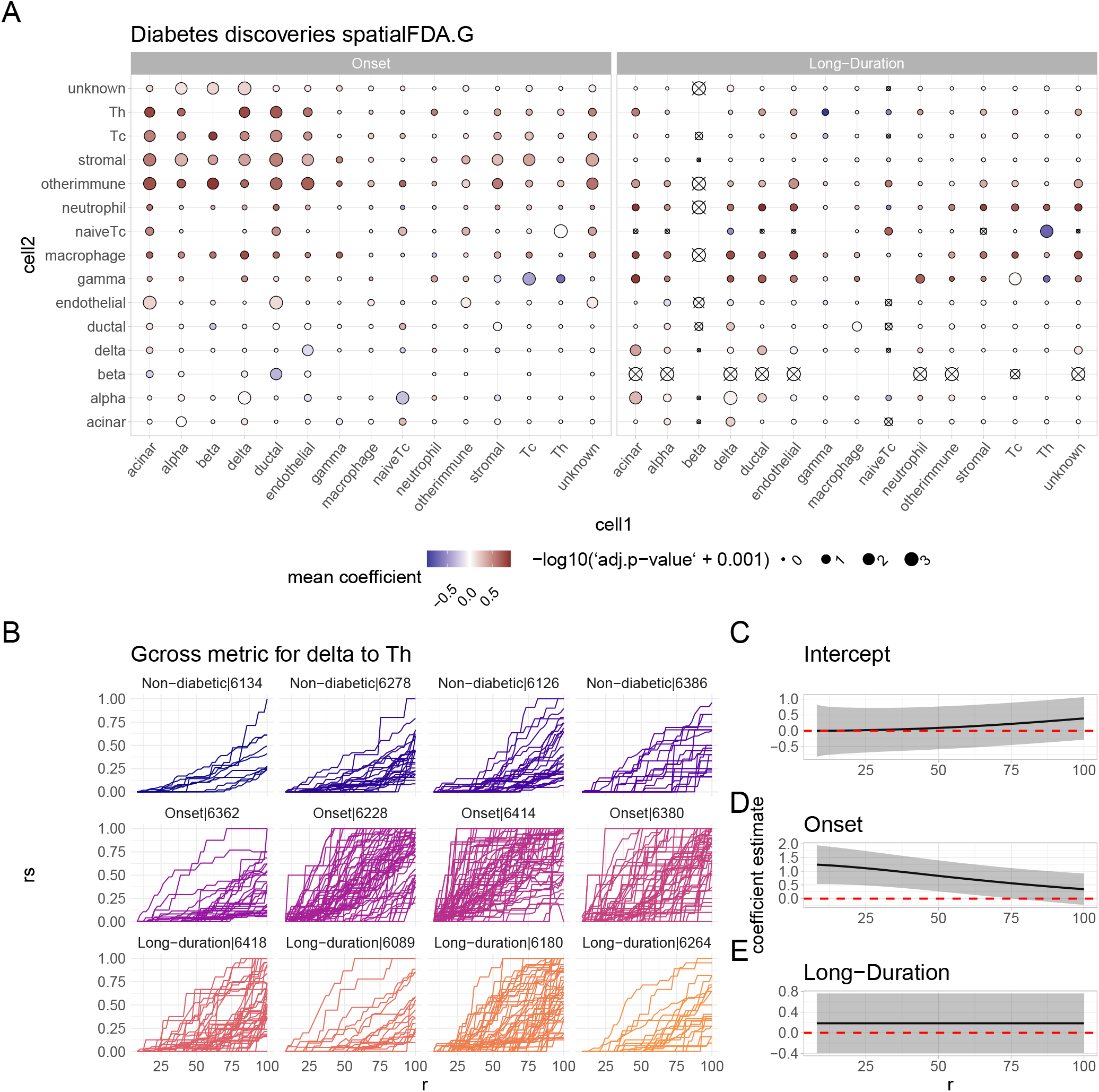
Results of the type-1 diabetes (T1D) CCoL analysis from Damond et al. **A** Bubble plot summary of the fGAMM results on *G* functions. The columns are the index cell types from which the *G* function is calculated and the rows are the cell types which are quantified in the spatially varying neighbourhood of the index cell type. The colour indicates the mean functional coefficient over the entire domain *r* whereas the size of the circles is the negative log *p*-values adjusted for multiple testing (FDR correction [51]). The two facets are the two T1D stages compared against non-diabetic samples. Combinations that fall below a minimum user-defined intensity threshold of 1*e* − 5 over all images are not considered and crossed through. **B** *G* functions for the cell type CCoL from *δ* to T helper cells. The *y*-axis is a reduced-sample estimator to account for edge effects [17, p. 285]. **C-E** Parameter estimates of functional coefficients. The intercept curve (C) estimates the mean curve of the reference (non-diabetic) category whereas D-E are the differences relative to the reference (non-diabetic) curve indicated by the red dashed line.

The intensity baseline `intensity.MM` finds no differences in the `spicyR` simulations (Figure S9). This verifies that only the local positions of the cells were perturbed in this simulation setting while the overall cell type proportions remained unchanged. The separation of cell type proportion and differential co-localisation effects was not directly possible in the Baker et al. simulation framework. This shows that the methods for differential CCoL can detect local re-arrangements that are independent of the overall cell type proportions.

We notice that there is no difference in the performance of mixed model variants and their corresponding fixed model alternatives, most likely due to the simulation not creating any sample-specific variation. Furthermore, we notice that the performance and calibration of `spicyR` (with an *L* function metric) become better if the simulated scale effect is in the middle of the integration range, whereas for `smoppix` (with a *G* function metric) both performance and calibration get worse with increasing radii. `spatialFDA.G` performs best when interactions occur at short scales whereas performance drops for changes beyond 50*µm*. Interestingly, this drop in TPR for larger radii in `spatialFDA.G` did not lead to an increase in FDR, meaning that `spatialFDA.G` is not miscalibrated for larger-scale interactions. `spatialFDA.L` shows a very stable performance across the simulated scale effects. The differences between *L* and *G* functions are expected theoretically, since they are sensitive to different aspects of the underlying point process: *G* functions summarise short-scale interactions while *L* functions summarise large-scale interactions [17, p. 295]. `SpaceANOVA` is well calibrated for close-by interactions (lambda 10-70) but shows worse calibration for interactions further away. Overall, `mxfda` without mixed-effects performs well in this simulation study, whereas the mixed-model led again to no detections.

Taken together, the simulations with varying length scales show that `spatialFDA.L` is sensitive to differential CCoL across length scales and `spatialFDA.G` performs well for short-scale interactions while maintaining a calibrated FDR for all scenarios. `mxfda` performs very well in this comparison, highlighting its good performance in scenarios without strong sample effects. The performance of the `spicyR` variants, `smoppix` and `SpaceANOVA` depends on correctly choosing an appropriate range for the interaction (not too wide and not too short).

### 2.4 Runtime comparison

Using the Canete et al. simulations, we performed a small runtime comparison in Figure S10. We ran all methods ten times and report the average elapsed time in seconds. In Figure S10A we report run-time as a function of the number of images in the dataset. Both `spatialFDA` and `mxfda` are among the fastest methods, followed by `spicyR`. `smoppix` and `SpaceANOVA` took substantially longer to compute. For `smoppix`, this is mainly due to the computation of the null model of the background cells whereas `SpaceANOVA` performs permutations for envelope-based summary functions. Both `smoppix` and `SpaceANOVA` offer fast implementations but do not result in strong sensitivity or specificity in our tests. Figure S10B shows the runtime as a function of the number of independent samples which are accounted for in the mixed-effects models as a random intercept. We note that `spatialFDA` scales rather poorly with the number of mixed-effects terms, a known limitation of random intercepts in the `mgcv` package that lies underneath `spatialFDA` [48]. An option in `spatialFDA` is to fit the models with `gamm4` instead, which uses efficient mixed-effects specifications of `lme4` [49, 28].

### 2.5 spatialFDA recovers spatial interactions in type-1 diabetes

To highlight the potential of `spatialFDA` to discover novel biological insights, we applied `spatialFDA.G` to a spatial proteomics dataset acquired with imaging mass cytometry (IMC) of type-1 diabetes (T1D) [5, 50]. Figure S11 shows three example FOVs for each of the three conditions, highlighting the strong structural differences in the disease progression of T1D. The choice of `spatialFDA.G` was due to our interest in close-range interactions (≤ 50*µm*) of pancreatic islets. T1D is an autoimmune disease that is characterised by the progressive destruction of insulin-producing *β* cells in the pancreas, often accompanied by a recruitment of cytotoxic and helper T cells early in the disease progression followed by a dispersion later in the disease [50]. It is thought that these immune cells play a role in the destruction of *β* cells early on in the disease [50]. To test for differences in CCoL across all cell types along T1D progression, we compared *non-diabetic* human donors with *onset* and *long-duration* diabetes (4 patients in each group). Each human donor sample in the dataset contains multiple FOVs resulting in a total of 845 images from 12 human donors.

To get an overview of the general CCoL of different cell types, we summarised the pairwise CCoL analysis across the disease progression in Figure 4A. We note that there are no combinations with *β* cells that passed the intensity threshold in long-duration diabetes, simply because there are few *β* cells left in long-duration T1D patients. Just by considering the intensity threshold in Figure 4A, we see the clear pattern of *β* cell destruction without having to look at the spatial metrics in detail.

Apart from the progressive loss of *β* cells in the course of T1D, the other striking change highlighted in Figure 4A is the increased CCoL of immune cells with the pancreatic islet cell types (*α, β* and *δ* cells) in onset T1D, as already described in Damond et al. (2019). We considered the CCoL of one combination in more detail, *δ*-Th cells. Figure 4B summarises the raw *G* function metrics across the (12 donor) samples. The raw *G* function metrics are non-smooth curves, which suggests that few Th cells were observed around *δ* cells. Figures 4C-E show the *functional* coefficients across scales of our statistical model; the intercept function (Figure 4C) represents the smoothed average *G* function across all non-diabetic FOVs. Figure 4D and Figure 4E show functions for *differential* CCoL to the onset diabetes and long-duration patients, respectively. A positive deviation from the red dashed reference line suggests stronger CCoL of *δ*-Th cells in the condition than in non-diabetic reference FOVs whereas a negative deviation means less relative CCoL. We note that there is no difference between long-duration and non-diabetic T1D functions (Figure 4D). However, there is an increased CCoL of Th and islet *δ* cells in onset T1D relative to non-diabetic functions (Figure 4E). Looking at the functional coefficients in Figure 4D in more detail, we noticed that there is a decrease in the function, suggesting that the increased CCoL is most pronounced close to the islets (10-50 *µ*m from *δ* cells).

An important step in statistical inference is checking model diagnostics. We considered both the adjusted R-squared for the entire model and the residual standard errors per condition in Figure S13. These two metrics give a broad overview of the confidence in the estimations. Notably, there is considerable variability in the adjusted R-squared (Figure S13A) across cell type combinations. For the pair *δ*-Th analysed above, the adjusted R-squared (0.6) and the Q-Q plot in Figure S14A suggest that the model is not capturing all variation in the curves. In addition, Figure S14B shows the large autocorrelation of residuals along the domain *r*. We use cluster-corrected sandwich covariance matrices in the statistical inference to make the results more robust against autocorrelated and variance heterogeneous residuals and to improve the calibration of the inference. However, the interpretation of specific point estimates according to the functional coefficients (Figure 4D-E) needs to be made with care and should rely on bootstrapped estimates instead [39, 40, 41]. In addition, we performed a leave-one-patient-out (LOO) stability analysis of the diabetes analysis. Figure S15A summarises the Jaccard similarity of significant results of the model on the full dataset with the LOO models without one patient respectively. We note that some patients are very influential for the significant results (patients 6228, 6126, 6414 with Jaccard similarity *<* 0.6). Figure S15B shows the concordance rate of each cell type combination across the LOO folds. The discussed *δ*-Th interaction is only significant in 50% of folds, highlighting considerable variability between patients. Figure S15C shows the stability of the mean effect size over the domain *r*. Most effect size estimates are relatively stable over the LOO folds. For both Figures S15B and S15C we notice that only few combinations are significant estimable for all folds in long-duration diabetes.

As we saw in the simulation studies, changes in differential CCoL can be convoluted with differences in intensities. To assess differences in intensities between the T1D stages, we plotted the average intensity per image across all cell types in Figure S12. Just looking at the average intensity per cell type for each image gives us a lot of information on T1D progression. For instance, we see the progressive loss of *β* cells in Figure S12. Other cell types such as *δ* cells show no obvious changes in intensity but were identified as significantly differentially co-localised in Figure 4. This highlights again that intensity and CCoL are distinct but non-exclusive concepts in spatial analysis.

We performed the same analysis with an *L* function instead of the *G* function above (Figure S16). We note that there are almost no significant differential co-localisations found across cell types for `spatialFDA.L`. This points to the biological process being on a smaller scale and less general, a setting that favours *G* functions. Since we also have large differences in overall intensity between cell types (Figure S12), we cannot exclude the possibility that *G* functions are sensitive to these intensity differences, despite our intensity adjustments (see Methods 5).

In order to put the findings of the diabetes dataset in relation to competitor methods, we ran the two methods that showed the best calibration in Figure 2, `spicyR` and `smoppix`, on the same dataset. This is an interesting comparison as both are simpler non-functional models that compare a scalar summary of a *L* and *G* functions (area between curves) between conditions. As we see in Figure S17 both `spicyR` and `smoppix` find fewer significant differential co-localisation cell type combinations than `spatialFDA.G`. Interestingly, the known biological immune recruitment during onset diabetes is not significant in either method. Since we do not have access to a ground truth annotation of this dataset, these comparisons remain at a qualitative level and do not let us make inferences about the individual false discovery rates.

### 2.6 spatialFDA relates changes in fibroblast associations in non-small cell lung cancer to the disease trajectory

As a second case study, we analysed the non-small cell lung cancer (NSCLC) IMC cohort from Cords et al. of 1071 patients and 2053 images (on average 2 images/cores per patient). Cords et al. focused their analysis on cancer-associated fibroblasts and described their importance in NSCLC. With `spatialFDA` we bring the added analysis of CCoL differences across the disease trajectory, an analysis that was not performed in Cords et al. Stage I was taken as reference and compared to stages II, III and IV for differential co-localisation of cells. Figure 5A shows example cores chosen at random from the dataset. We notice a very unstructured pattern in stage IV in comparison to the other stages. Figure 5B summarises the pairwise differential CCoL across the disease trajectory. In general, we find only a few significant CCoL differences at the 5% FDR level in this dataset and most interactions are more spaced in later stages than the stage I reference. The largest number of interaction differences is found for tumour, vessel and fibroblast cells. In Figure 5C, we look in more detail at the tumour-fibroblast interactions. We notice that there are fewer fibroblasts found in the vicinity of tumour cells in stages II and III in comparison to stage I, but there is no difference relative to stage IV. This pattern is also visible (though not significant) in Figure 5D for the vessel-fibroblast interactions.

**Figure 5:**
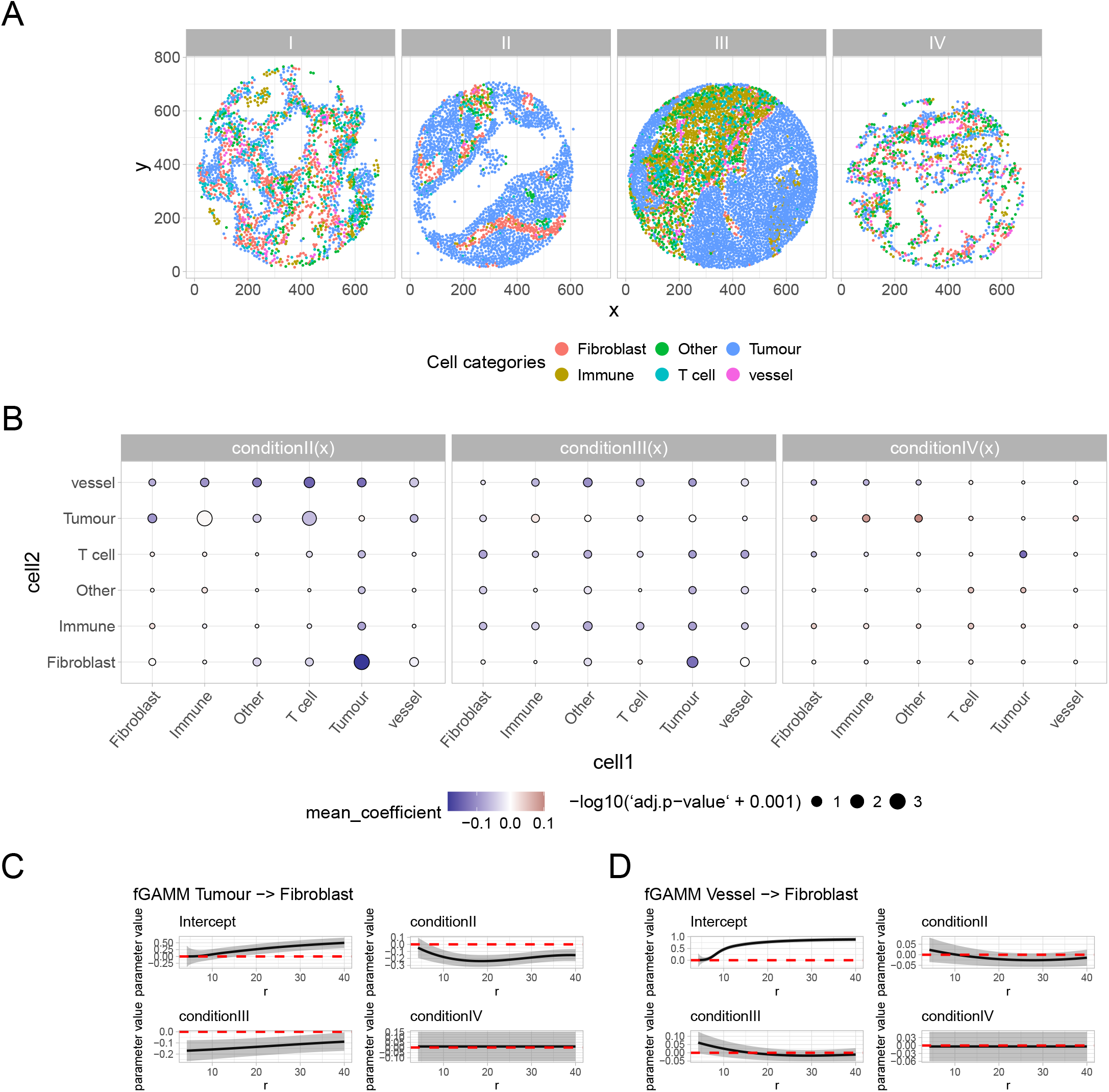
Results from the differential CCoL analysis of the non-small cell lung cancer (NSCLC) study from Cords et al. **A** Example IMC cores across the disease stages I-IV coloured by the broad cell type categories considered in the CCoL analysis. **B** Heatmap of the pairwise CCoL results. The colour indicates the mean functional coefficient over the entire domain *r* whereas the size of the circles is the negative log *p*-values adjusted for multiple testing (FDR correction [51]). **C-D** functional coefficients across the domain *r* for the interactions Tumour-Fibroblast and Vessel-Fibroblast

The experimental design of this study is a challenge for `spatialFDA` in two ways: i) fitting mixed-effects models with *>* 1000 random intercepts becomes computationally expensive in `mgcv` and ii) estimating random intercepts from an average of two observations is challenging in terms of identifiability. A reasonable alternative is to randomly choose one observation/image per patient and compare 1071 identically and independently distributed samples with a fixed-effects model instead. We show the results of this fixed-effects model in Figure S18, noting that there are only minor differences to the mixed-effects model on the full dataset (Figure 5).

In summary, we have investigated the key spatial interactions surrounding fibroblasts from the Cords et al. study. In addition to their findings, we find clear fibroblast-tumour differences across stages in CCoL that were not presented in the original manuscript. We also highlight the effect of the study’s experimental design on modeling choices (i.e., fixed versus mixed modeling).

## 3 Discussion

We presented the novel method `spatialFDA` to perform spatial CCoL analyses across scales. `spatialFDA` is a free and open-source `Bioconductor` package that can be easily integrated with other spatial omics analysis packages. We have shown that `spatialFDA` performs well in controlled simulation scenarios, being sensitive across different simulation settings while controlling the false discovery rate. In a biological use case on type-1 diabetes, we uncovered known biological effects from previous studies about the spatial organisation across the diabetes disease progression [50]. In NSCLC, `spatialFDA` helped to find patterns of fibroblast associations with both tumour and vessel structures that are different across the disease stages, a finding that was not previously described in Cords et al.

In contrast to most methods for differential CCoL, `spatialFDA` maintains the spatial scale of the data throughout the analysis by providing model coefficients as a function of the radius *r*. At its core, `spatialFDA` computes an average spatial metric across one FOV. Of course, this can underestimate the within-FOV heterogeneity. An option to improve on this would be to not view the FOV as the unit for average spatial metric computation, but rather spatial clusters or histopathologically annotated regions.

A question that arises in the analysis of multi-sample spatial omics experiments is how to treat several layers of variability (e.g., repeated measurements). In the case study of T1D, we had data from three conditions with several samples per condition, and multiple FOVs per sample were acquired. The smallest independent unit was therefore the sample, and the FOVs have to be treated as non-independent. In `spatialFDA`, this can be readily addressed with generalised functional mixed-effects models, by flexibly accounting for different levels of variability [37, 36, 38]. In the case study on NSCLC, we had the other challenge of many independent patient samples and few replicates per patient. This is a difficult setup for mixed-effects models as random intercepts per patient are not easily identified with few observations (∼ 2 in the NSCLC study) and models with thousands of random intercepts scale poorly in general. Therefore, we advocate subsampling such datasets to have one observation per patient and fit a fixed-effects model instead. This flexibility in model definition should be guided with model diagnostics such as quantile-quantile and autocorrelation plots of the model residuals as well as general considerations on experimental design.

One challenge in the inference of spatial functions is their constrained nature. `spatialFDA` addresses this by correcting the covariance matrices post-hoc for correlations along the functional domain *r* with sandwich estimators. Future work should investigate the possibility to model these functional constraints directly in the statistical model.

One general limitation of all CCoL methods considered is the approximation of cells as points in space. This neglects the natural spacing of cells due to the cell body. We correct our inference for the average nearest-neighbour distance between cells. This is only a reasonable correction if cells have uniform and round shapes.

Another assumption at the foundation of all CCoL methods is that the underlying spatial patterns are invariant to rotations (isotropy). This is an unrealistic simplification since many biological tissues are comprised of anisotropic patterns. Anisotropic spatial statistics is an open field of research and may offer important tools in the future for biological tissues [53, 54].

Our simulation study, while able to better mimic densely-packed architectures observed in real tissue, also carries the caveat that cell type proportion changes are introduced simultaneously to cell type interaction changes. Fundamentally, this is not a disadvantage but depends on the definition of CCoL and highlights that these two aspects are not always fully separable. The reduction of a cell type, for example, cells being pushed outside of the FOV, can be a form of (unobserved) CCoL. We argue that both intensity/proportion and CCoL differences of all cell types should be studied collectively in spatial omics data. Furthermore, choices for *K* and *L* functions over *G* functions in the analysis will separate the two effects better.

In addition, we did not explicitly simulate data that violate the assumptions of CCoL methods to check for robustness towards misspecifications [55]. The main untested assumption of all CCoL methods is first- and second-order stationarity of the underlying point pattern [17].

## 4 Conclusion

We have introduced `spatialFDA`, a novel method for differential CCoL analysis. In two simulation studies, we have shown that `spatialFDA` is a sensitive method for differential CCoL in controlled settings, controlling error rates better than most competitor approaches. In a biological study of T1D, `spatialFDA` uncovered known spatial effects with respect to the disease biology, while retaining information on the spatial scale of these interactions.

## 5 Methods

### 5.1 spatialFDA methodology

#### 5.1.1 Quantification of spatial data with spatial metrics

Spatial arrangements are summarised with spatial metrics. These functions have in common that the domain consists of increasing radii *r*. `spatialFDA` takes a sequence of radii *r* and a cell type (pair) as input and computes function values implemented in the `R` package `spatstat.explore` as output [44].

One challenge in spatial metric calculation is how to account for apparent inhomogeneity due to technical artefacts such as missing tissue regions. In `spatialFDA`, researchers can pass a custom observation window restricted to high-density cell regions to the metrics calculation [24].

The two main categories of spatial metrics we consider in `spatialFDA` are correlation and spacing functions for discrete cell type marks.

##### Correlation

Spatial correlation describes the dependence of points on each other. The null model for most spatial statistics is a Poisson process where points are independent of each other. The three main types of interactions one can distinguish are i) regularity (functions lie below the Poisson distribution), ii) independence (functions follow Poisson distribution) and iii) clustering (functions lie above Poisson distribution) [17, p. 199].

A common technique to assess spatial correlation is via Ripley’s *K* function [17, p. 203 ff.] [22, 21]. The empirical Ripley’s *K* function is defined as:

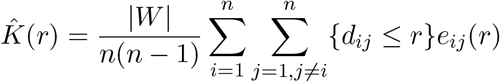

where |*W* | describes the window size of the field of view, *n* the number of points in the window, *d*_*ij*_ the distance between point *i* and *j* and *e*_*ij*_(*r*) an edge correction measure [17, p. 204][22, 21].

The correction of the window size |*W* |, the number of points *n* and the edge correction *e*_*ij*_(*r*) allow for a comparison of fields of view with differing sizes and numbers of points [17, p. 204].

There are many variants of Ripley’s *K* that compare univariate, bivariate or even multivariate patterns, correct for inhomogeneous point patterns or different scale effects [17, p. 203 ff.].

Most often, Ripley’s *K* function is variance stabilised, which in turn is called Besag’s *L* function [23, 56].

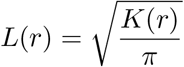

##### Spacing

The other main class of spatial metrics is based on the concept of spacing. These summary statistics are complementary to the correlation summaries described above. Spacing functions are based on distances between points. These can either be pairwise distances, nearest-neighbour distances or empty-space distances [17, p. 255].

The nearest-neighbour distance distribution function *G*(*r*) is based on nearest-neighbour distances *d*_*i*_ = min_*j* ≠*i*_ ||*x*_*j*_ − *x*_*i*_||. This can also be written as *d*_*i*_ = *d*(*x*_*i*_, **x** \ *x*_*i*_). For any location, *u* in a stationary point process we then obtain the distance *d*(*u*, **X** \ *u*). The cumulative distribution function of the nearest-neighbour distance is then [17, p. 262]:

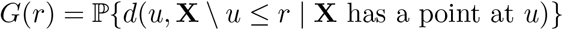

In the completely spatially random case the nearest-neighbour function *G*(*r*) is given by [17, p. 264]

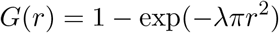

#### 5.1.2 Inference on spatial metrics with functional GAMMs (fGAMMs)

To test for associations between sample and condition covariates and the spatial metrics, we use a fGAMM implemented in `functionalGAM`. This function is a wrapper around the `refund` function `pffr` [37, 36, 45]. In addition, it allows for the specification of the experimental design with a model matrix and a model formula given a `SpatialExperiment` object [43].

The model we use is defined as

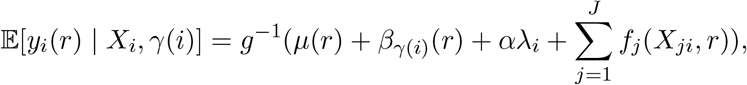

where *y*_*i*_(*r*) are the functional response functions (here the spatial metrics), *g* is an optional link function, *µ*(*r*) is the global intercept function (for the reference category), *β*_*γ*(*i*)_(*r*) is the functional random intercept per grouping variable *γ*(*i*), and *f*_*j*_(*X*_*ji*_, *r*) are the (linear/unpenalised or smooth/penalised) contributions of the other scalar covariates to the additive predictor. The term *αλ*_*i*_ is an image-specific intensity covariate adjustment factor for the query cell type (in a cross function *A* → *B* it will adjust for cell type *B*). This is important for *G* functions which are not already intensity corrected like *K* or *L* functions. The intensity adjustment feature is only implemented for *G* functions as a default in `spatialFDA`. Except for Figure S7, all `spatialFDA.G` versions are intensity adjusted in the model, whereas for `spatialFDA.L` or `spatialFDA.K` the functional outcomes are intrinsically intensity-adjusted by construction.

In the case of `spatialFDA` and the associated function `spatialInference`, we model functions with a Gaussian distribution and a log link to ensure positivity of the functional response along the domain *r*. Other potentially valid options provided by our software implementation such as Beta-distributions for *G* functions or Gamma-distributions for *L* functions did not converge during fitting or gave worse AIC results.

Inference of spatial metrics with fGAMM can be affected by both autocorrelation and heteroscedasticity of the residuals along the domain *r*. Functional data derived from spatial omics data frequently violate the underlying assumption of independently and identically distributed (Gaussian) residuals of these models. To obtain covariance estimates that are more robust to both autocorrelation and heteroscedasticity, we have implemented cluster-robust covariance matrix estimators (*V*_*CL*_) via sandwich correction. These estimators work by allowing arbitrary within-function correlation and heteroscedasticity of the residuals while only requiring that the individual functional responses are independent.

The cluster-robust sandwich covariance estimators are defined as

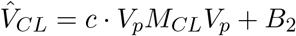

where *c* = *m/*(*m*−1) is a CR1-type finite-sample adjustment, *V*_*p*_ ∈ ℝ^*p*×*p*^ the posterior Bayesian covariance matrix (so-called bread of the sandwich), 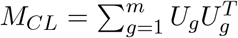 where *U*_*g*_ are the aggregated cluster-level score functions (in this case *g* are the functions) (so-called meat of the sandwich) and *B*_2_ = *V*_*p*_ −*V*_*e*_ ∈ ℝ^*p*×*p*^, (*V*_*e*_ ∈ ℝ^*p*×*p*^ is the frequentist covariance matrix) for a Bayesian covariance correction or *B*_2_ = 0 in the frequentist case [39, 40, 41].

Since cells have a diameter and do not overlap, all raw functions have a constant zero part at the beginning. Because this leads to difficulties in fitting mean and variance, we remove the constant all-zero part of the functions. We estimate the cell spacing due to the cytoplasm via the weighted mean of the minimum nearest-neighbour distance per image and filter any radii *r* smaller than this cut-off.

Statistical significance over the entire domain is assessed via a global *F* -test. The *F* -test allows us to test hypotheses on the entire function, i.e., whether the function is different from the reference function at any radius *r*. Specifically, the *F* -statistic tests the hypotheses

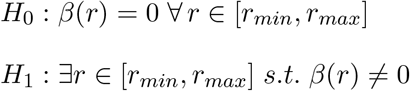

where *r*_*min*_ is the average nearest-neighbour corrected centroid distance between cells (see above) and *r*_*max*_ is the user-provided maximum search radius. `spatialFDA` provides a heuristic for the maximum search radius by computing the minimum bounding box for each image across the dataset. The maximum search radius heuristic is then half of the smallest minimum bounding box in the dataset.

This *F* -test for “no effect” uses a cluster-robust covariance matrix which corrects for heteroskedasticity and autocorrelation across *r*.

Similarly, the confidence bands of the coefficient functions are corrected for autocorrelation and heteroscedasticity of the residual errors, but since they are only pointwise and approximate, they should be interpreted with care. To obtain a more reliable indication of the significance at a specific radius *r*, a bootstrapped confidence band should be calculated [57, 58], especially in smaller samples.

#### 5.1.3 Heatmap of pairwise combinations

The fGAMM can be calculated over a range of *n* cell type combinations in a pairwise manner. To generate an overview of these *n*×*n* tests we generate a heatmap. Per test pair we summarise the significance as the *p*-value of the *F* -test adjusted for multiple testing with FDR correction [51]. The functional coefficient is summarised as the mean coefficient over the functional domain *r*. This allows us to obtain a picture of the effect along with the directionality of the interaction in comparison to the intercept group, in addition to a notion of its statistical significance from the *F* -test.

### 5.2 Simulation framework

The simulations are extended from the published simulation framework of Baker et al. (2023).

In a first step, an empty tissue scaffold is created with random circle packing. This works by placing circles in a defined window (1000 × 1000) while controlling the minimum and maximum circle radius *r* ∈ [5, 20]. From the image, a graph is constructed with the nodes being the centroids of the circles. Undirected edges are placed between the centroids of adjacent circles.

The cell typing takes two arguments: i) *p* ∈ ℝ^*K*^ the vector of the *K* cell type proportions in the scaffold and ii) *H* ∈ ℝ^*K*×*K*^ the matrix of cell type interaction probabilities. The cell type proportion for cell type *i* (*p*_*i*_) indicates the probability of finding this cell type in the tissue scaffold. The interaction probability of cell types *i* and *j* (*H*_*i,j*_) parametrises how likely it is to find cell type *j* adjacent to cell type *i* on the constructed graph.

To create a multi-condition, multi-sample framework, one cell type interaction probability is sampled from *h*_00_ ∼ *N* (*µ*_*condition*_ + *δ*_*sample*_, *σ*) where *µ*_*condition*_ is a condition-specific mean, *δ*_*sample*_ is a sample-specific offset and *σ* an equal variance parameter between samples. Between conditions we parametrize *µ*_*control*_ = 0.5 and *µ*_*perturbed*_ = {0.3, 0.2}, whereas between samples the offset varies according to *δ*_*sample*_ ∼ *U*(−0.2, 0.2). The variance *σ* = 0.25 is fixed. To obtain valid probabilities for *h*_00_ ∈ [0, 1], we truncate the Normal distribution *N* (*µ*_*condition*_ + *δ*_*sample*_, *σ*) with symmetric rejection sampling. The probability vector *p* is then row-normalised and the matrix *H* is row- and column-normalised to obtain valid probabilities summing to one.

The cell types are then heuristically assigned in two steps. A start node *ν*_*i*_ is selected from the graph and the cell type of this starting node *ν*_*i*_ is assigned according to 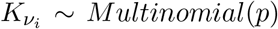. The probabilities of the *n* neighbours of *ν*_*i*_ are chosen according to *ν*_*n*_ ∼ *Multinomial*(*H*_*K*∗_) [47].

The cell typing is repeated to obtain several varying annotated tissue sections per sample. In addition, a smaller field of view (500 × 500) is chosen at random from the larger section to mimic the selection of regions of interest from a tissue section. With this repetition of cell typing, we model variation within a sample.

We run the CCoL methods on these simulated fields of view. By repeating the simulation and the inference with the methods 500 times, we obtain a notion of power of each of the methods.

In order to create more simulation scenarios we change the cell type proportion vector (*p*_0_ = {0.15, 0.25, 0.6}).

### 5.3 Canete et al. simulation framework

The second simulation is taken from Canete et al. (2022) and adapted to accommodate both `spicyR` and `spatialFDA`. They simulate 40 patients across two conditions. Each patient has 3 images, each containing 2 cell types *A* and *B*. First, a random Poisson process for cell type *A* with rate *λ*_*A*_ ∼ [20, 40, …, 400] is simulated. The densities of cells of type *A* are simulated as a disc kernel of radius *σ*. The radius of the disc changes between simulation scenarios *σ* ∈ [10, 20, 30, …, 100]. The cells of type *B* are then simulated from a Poisson process with rate *λ*_*B*_ ∼ [20, 40, …, 400] multiplied by the density of the kernel *σ* for the control group and the density *σ* + *σ/*2 for the cases. More details can be found in the original publication [29].

### 5.4 Performance evaluation in simulation study

For the Baker et al. simulation we assign true negatives as our simulated controls (no change relative to the reference group) and true positives as our simulated perturbations (20% change relative to controls). We mix true positives and true negatives at equal proportions. In addition to the simulated ground truth, we provide the *p*-values of those simulations. For the Canete et al. simulation, we calculate true positives and true negatives for each simulation setting. For both simulations, we calculate both true-positive rate vs. achieved false-discovery rate and receiver operating characteristic curves using the `iCOBRA` package [59]. Monte Carlo standard errors are computed as 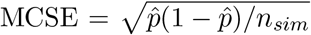, where 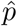 is the observed FDR of the method, and are used to construct Wald-type confidence intervals [55].

### 5.5 Scalar methods

The so-called scalar methods summarise the information of the spatial metrics into a single number and compare these scalars with linear (mixed) models. These include our baseline intensity mixed model, `spicyR` and `smoppix` [29, 30]. In the intensity mixed model we compute the average homogeneous intensity of the perturbed cell type *x* as 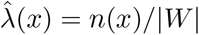 where |*W* | is the area of the field of view. This gives one scalar per image which we compare between controls and cases with a linear mixed model with a random effect per sample. In the Canete et al. simulation the `intensity.MM` baseline is adapted to a cross-cell type comparison (*B* → *A*). We compute the difference in average intensity for both cell types *A* and *B* separately, as both could influence the cross CCoL. We report the smaller of the two *p*-values for both models in the performance evaluation.

`spicyR` is a published method with different modes of computing the summary of the *L* function per image. We compute the area between the estimated *L* function and the theoretical Poisson curve as 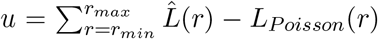. This summary is compared with either a linear or linear mixed model [29].

`smoppix` is another method that compares the ABC of *G* functions. The *G* functions of individual cells *g* and *h* are summarised by the ABC and are aggregated per point pattern 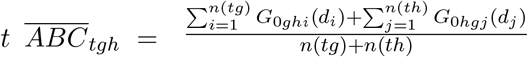 where *G*_0*ghi*_ is the null distribution of feature *h* to cell *i* of feature *g*. The null distribution is estimated from the background of all cells in the point pattern *t*. The aggregated ABC values per point pattern *t* are then compared using a linear mixed model [30].

### 5.6 Functional methods

The functional methods compare the entire function across its domain between conditions and samples. The first functional method in the comparison is `SpaceANOVA`. It uses two variants of a functional ANOVA. The first is the one-way functional ANOVA:

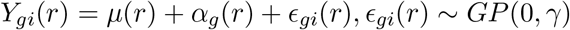

where *µ*(*r*) is the common, *α*_*g*_(*r*) the group-specific and *ϵ*_*gi*_(*r*) the subject-specific mean function. The statistical test is then formulated as

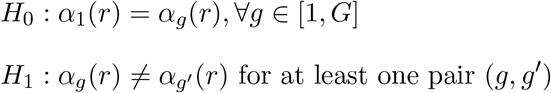

and tested with an *F* -test given by the formula:

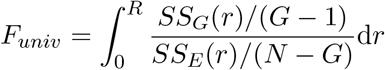

*SS*_*E*_ and *SS*_*G*_ stand for the within-group and the between-group sums of squares, *G* for the number of groups and *N* the total number of images.

The second option is the two-way functional ANOVA:

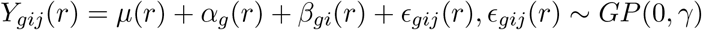

where *µ*(*r*) is the common, *α*_*g*_(*r*) the group-specific, *β*_*gi*_(*r*) the subject-specific, and *ϵ*_*gij*_(*r*) the image-specific mean function.

The hypotheses are the same as in the univariate functional ANOVA [32]. The *F* -test follows:

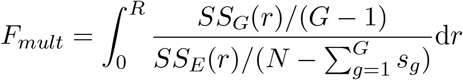

The other functional method is called `mxfda` [33]. Unlike all other methods mentioned before, the spatial metric is a covariate in this model instead of the response. The function can be chosen flexibly; the most common are *K* and *G* functions. `mxfda` offers modelling options for survival, binary and continuous responses. In our simulation study we use a functional logistic regression model

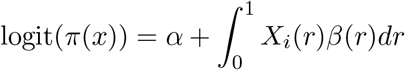

where *π*(*x*) = *P* (*Y* = 1|*X* = *x*) is a conditional probability, *α* is an intercept parameter, *X*_*i*_(*r*) is the functional predictor (*L* or *G* functions) and *β*(*r*) the coefficient function [60].

Significance of the functional coefficients is tested via a global *F* -test, similar to `spatialFDA` [58].

### 5.7 Diabetes data analysis

We reanalyse a published imaging mass cytometry dataset of type-1 diabetes [50]. We quantify differential CCoL of all cell type pairs across the disease progression. We compare non-diabetic patients as a reference group to onset and long-duration diabetes using `spatialFDA` with *G* and *L* functions.

In order to assess the stability of the `spatialFDA.G` results, we perform a leave-one-patient-out (LOO) stability analysis. We compare the full-dataset run to our LOO runs in terms of the per-patient Jaccard index, intersection over union of significant discoveries between full-dataset and LOO runs, the cell type pair concordance rate for each model term, defined as the proportion of LOO folds in which that term remained significant as well as the stability of the mean estimated coefficient over the functional domain *r* for each pair. For the cell type pair significance and effect size LOO analysis we restrict the analysis to pairs that are significant and that are being estimated in each LOO run.

In addition, we tested the two best calibrated competitor methods from the simulation study, `spicyR.MM` and `smoppix`, on the same diabetes dataset.

### 5.8 Non-small cell lung cancer data analysis

As a second biological use-case, we reanalyse a published imaging mass cytometry dataset of non-small cell lung cancer (NSCLC) [52]. Cords et al. provide three cell type classification levels. At the coarsest level, we quantify differential CCoL of all pairs across the disease progression. The reference group is stage I NSCLC and we compare stages II, III and IV respectively to the reference with `spatialFDA.G`.

## 6 Data availability

The raw data of the diabetes study can be accessed on Mendeley Data https://data.mendeley.com/datasets/cydmwsfztj/2 and are published under a CC-BY-4.0 license with the associated publication [50].

The raw data of the NSCLC study can be accessed on Zenodo https://zenodo.org/records/7961844 and are published under a CC-BY-4.0 license with the associated publication [52].

## 7 Code availability

`spatialFDA` has been a `Bioconductor` package since the 3.21 release. The source code can be accessed at https://github.com/mjemons/spatialFDA.

The entire analysis of this paper is bundled in a `snakemake` [61] workflow and available at https://github.com/mjemons/spatialfda-manuscript.

## 8 Acknowledgements

We thank Reinhard Furrer, Torsten Hothorn, Constantin Ahlmann-Eltze and Wolfgang Huber for helpful discussions on developing the method and comments on the manuscript. We thank Izaskun Mallona, Charlotte Soneson and David Wissel for their help with the simulations and performance evaluations.

## Artificial intelligence tools

We acknowledge the use of artificial intelligence tools in this work. Claude (Sonnet 5 and lower) and ChatGPT (5.6 and lower) were used for literature research, review paragraphs for consistency, check grammar and spelling, suggest alternative phrasing for individual sentences, check mathematical notation, assist in generating parts of the `spatialFDA` package and analysis code. All content was assessed manually and we take full responsibility for all content in this work.

## 9 Author contributions

**ME**: Methodology, Software, Writing - Original Draft **FS**: Methodology, Software, Writing - Review & Editing **SG**: Software, Writing - Review & Editing **EP**: Methodology, Writing - Review & Editing **MDR**: Conceptualization, Methodology, Writing - Review & Editing, Supervision, Funding acquisition

