## Supplementary Information for "Differential co-localisation analysis of multi-sample and multi-condition experiments with spatialFDA"

September 10, 2026

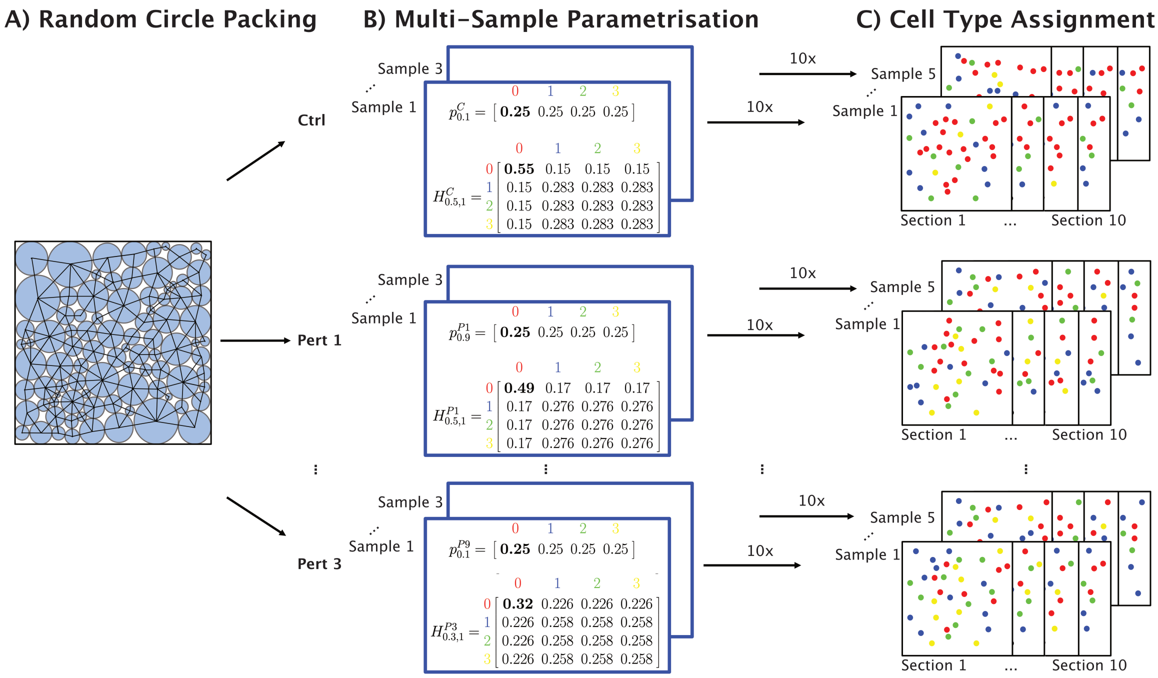


Figure S1: Figure summarising the simulation framework based on Baker et al. **A** The first step is random circle packing. For the different conditions (controls and all perturbations), the square observation window of 1000 x1000 µm is packed with random circles with radius $r \in [5, 20]$. Per condition this done for five samples resulting in five tissue scaffolds generated per sample. A graph is constructed on top of these circles with nodes at the circle centres and edges between touching circles. **B** The second step is the parametrisation for a multi-sample and multi-condition experiment. The cell type assignment requires two parameters, the cell type proportion vector $p$, and the cell type interaction matrix $H$. The cell type proportion vector $p$ changes for cell type 0 between simulation scenarios but is fixed for one power curve. The interaction probability matrix $H$ changes for the cell type 0. The interaction probability $h_{00}$ differs between the perturbations and is sampled from a $[0, 1]$ truncated Normal distribution with a condition specific mean and an equal standard deviation for all samples. This results in perturbation/sample specific interaction matrices $H_{pert,sample}$ with row- and column-normalised probabilities for which one example matrix of three is indicated in the two shown perturbations. **C** Shows the actual cell type assignment based on the tissue scaffolds from A) and the parameters from B). For each tissue scaffold, the centroids are assigned one of four cell type labels (0-3). This sampling is repeated ten times and results in 10 annotated cell type sections per sample, in total 50 sections per condition.

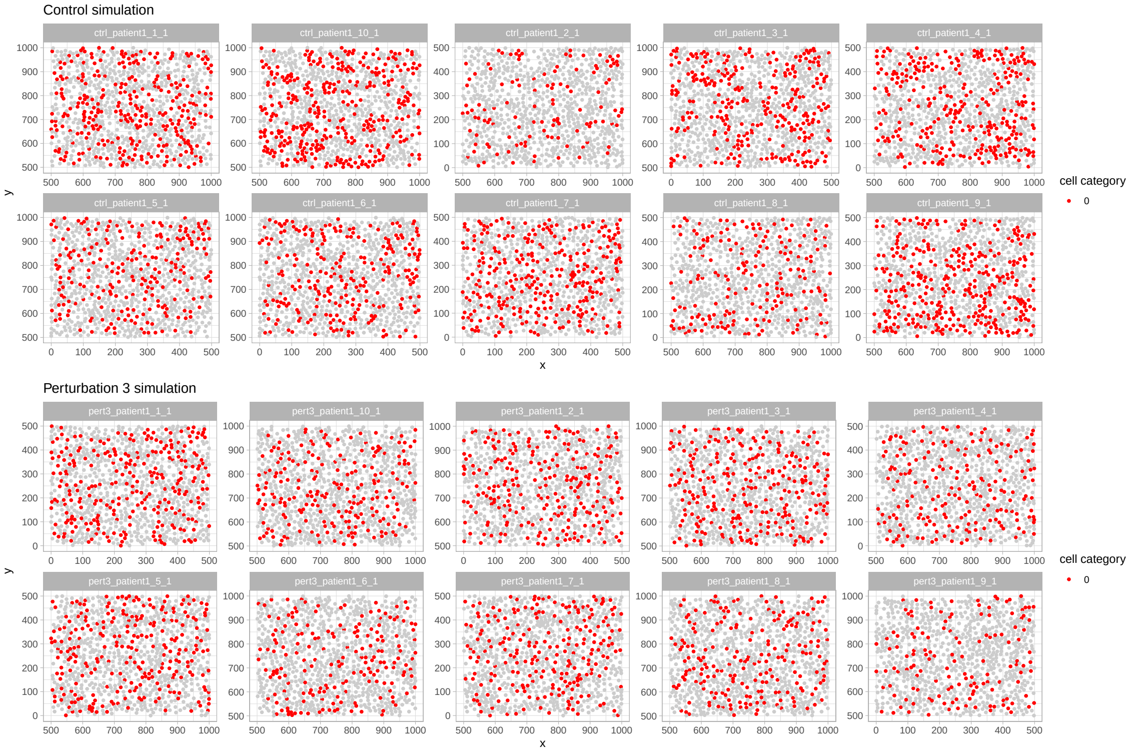


Figure S2: Simulated point patterns for two patients. Red indicates the cell type 0 that is perturbed, for visualisation all other cell types are coloured in grey. The first are control simulations with a cell type interaction probability $H_{0,0} = 0.5$. Clearly visible is the clustering but also the simulated within patient variance in the interaction probability. The second patient are perturbation simulations with $H_{0,0} = 0.3$. An apparent smaller degree of clustering of cell type 0 can be found but still a similar within patient heterogeneity in the interaction probability as in the control simulation.

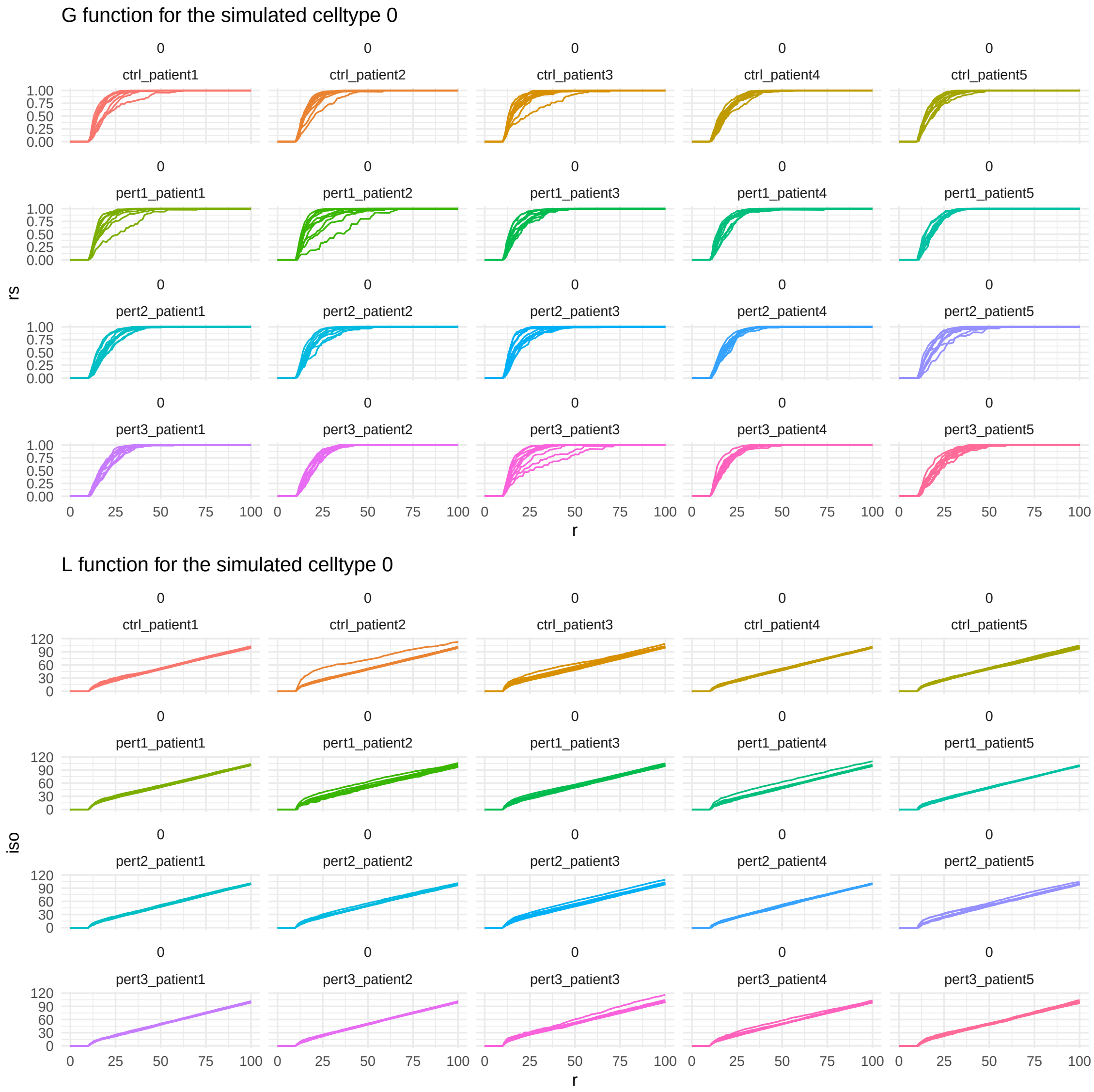


Figure S3: $G$ and $L$ functions for one representative simulation round in which the interaction probability for cell type 0 is changed. For each simulated condition the interaction probability $H_{0,0}$ changes (control = 0.5, perturbation 1 = 0.5, perturbation 2 = 0.4 and perturbation 3 = 0.3) and 5 patients are simulated with 10 images each.

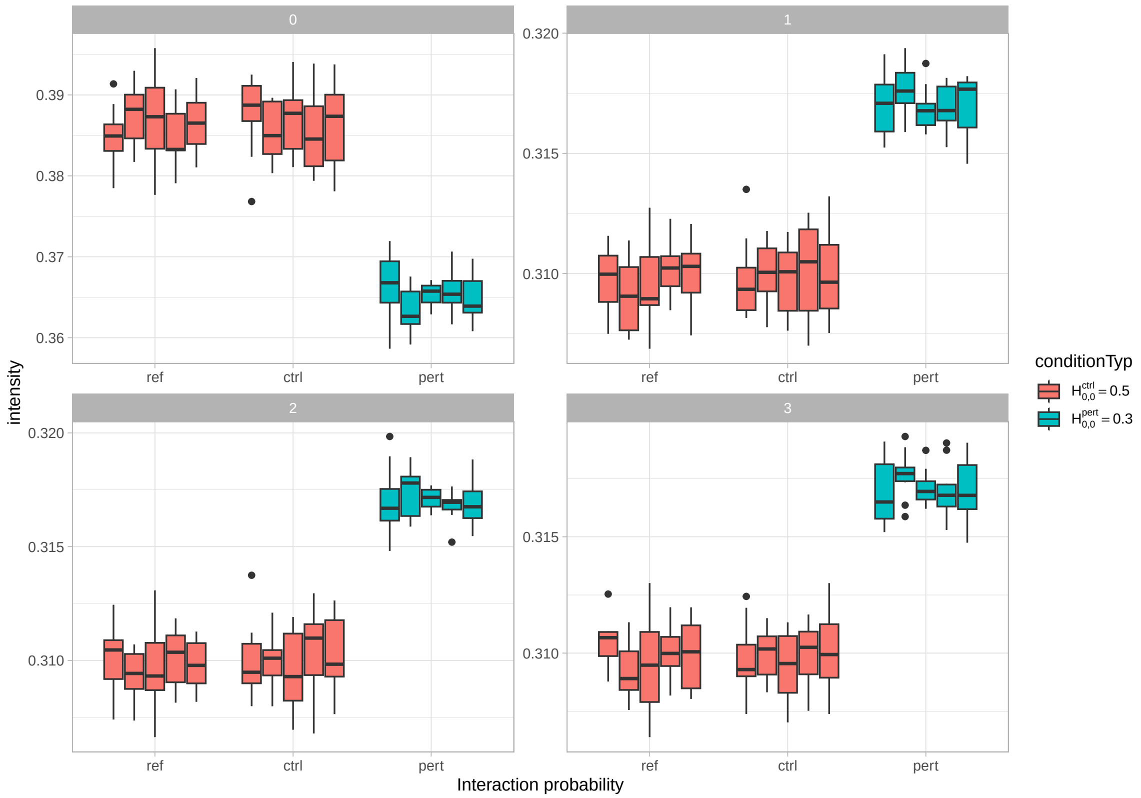


Figure S4: Boxplot of the marginal intensities per cell type after the simulation. The facets correspond to the cell types [0-3], the $y$-axis to the average intensity per FOV. The $x$-axis corresponds to the simulated interaction probabilities between conditions (ref $= H_{00} = 0.5$, ctrl $= H_{00} = 0.5$, pert $= H_{00} = 0.3$. For the target cell type 0, just changing the interaction probability $H_{00}$ already leads to a drop in the average intensity. For the other cell types, the reduction in $H_{00}$ leads to an increase in the average intensity per FOV.

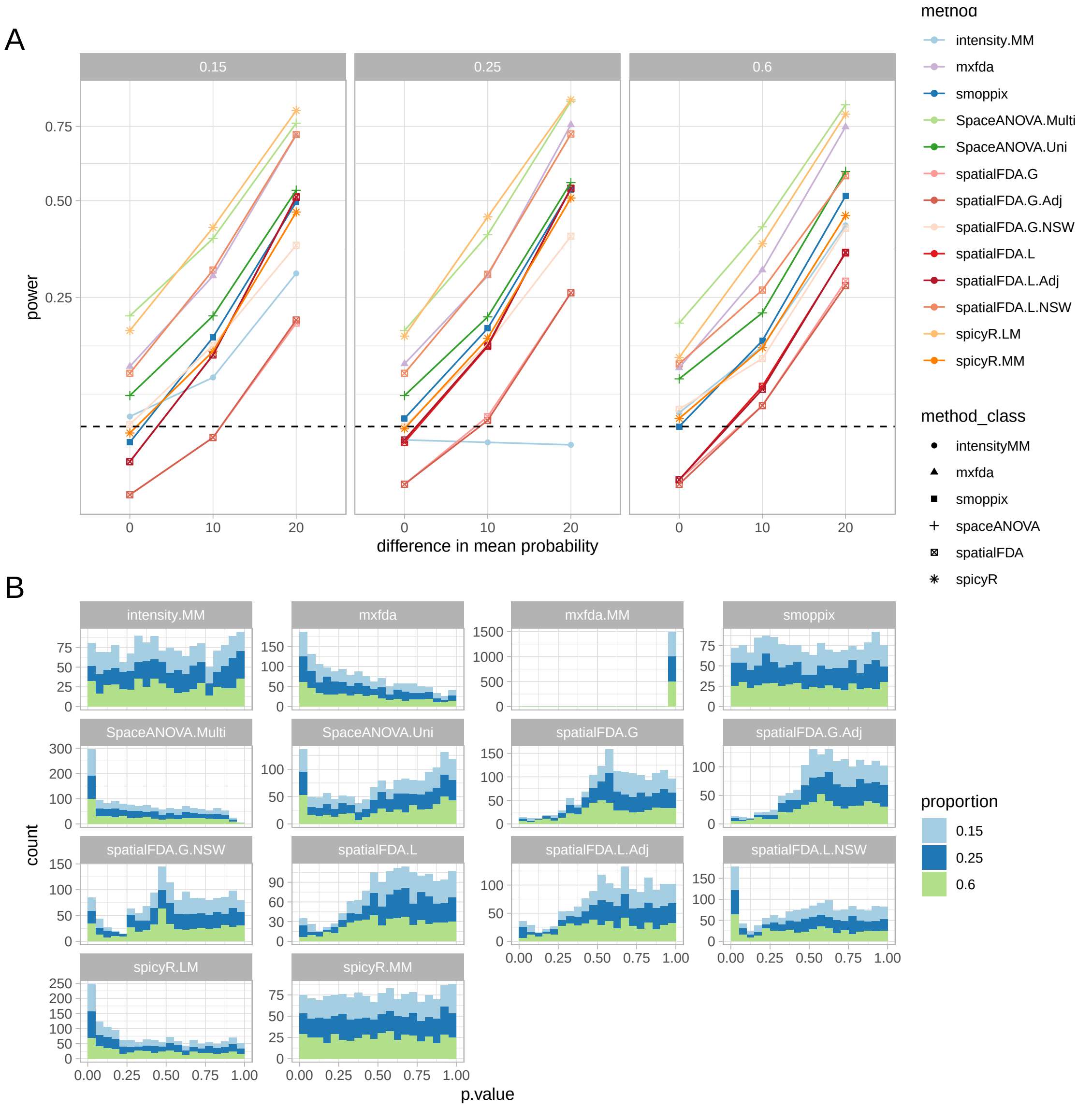


Figure S5: **A** Results of the simulation study adapted from Baker et al. The $x$-axis corresponds to the difference in mean probability between the controls and the perturbations. This means 0 is the null simulation of no difference and 20 corresponds to a difference of 20% between controls and perturbations. The $y$-axis is the power of the test (the percentage of positive results detected among all simulated scenarios). The shapes indicate the methods, whereas the colours indicate the method version outlined in table 1. The black dashed line indicates the positive rate at 0.05, which is what we would expect under the null hypothesis. The multiple plots are ordered by increasing prior cell type proportions. **B** $p$-value distribution under the null hypothesis. The colour indicate different prior cell type proportions of cell type 0 in the simulation.

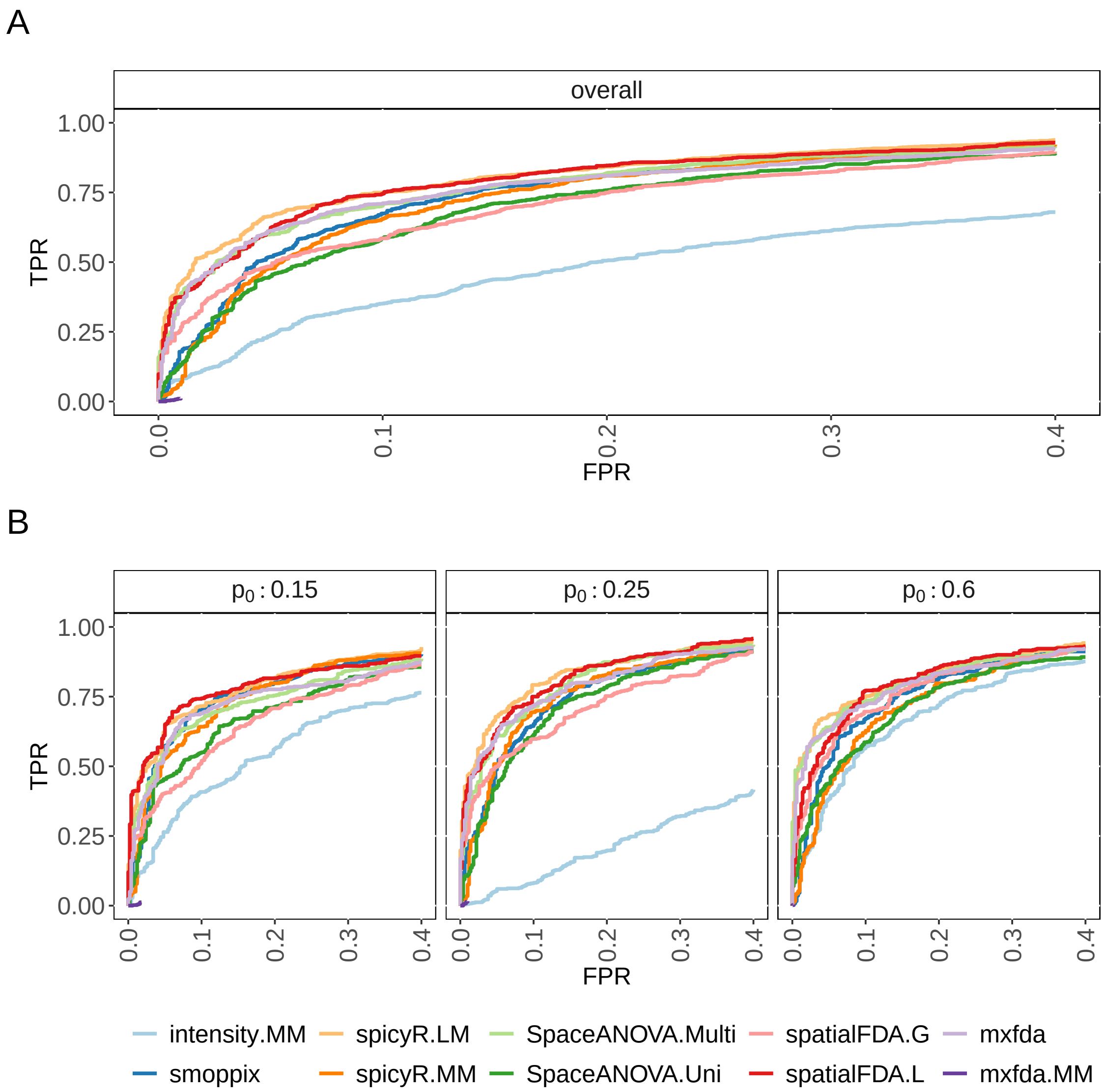


Figure S6: Receiver operating characteristics curve of true-positive rate vs. false-positive rate for all the different methods in the Baker et al. simulation. **A** Overall performance results across all simulations. **B** Performance results split by the prior cell type proportion of the perturbed cell type.

A

B

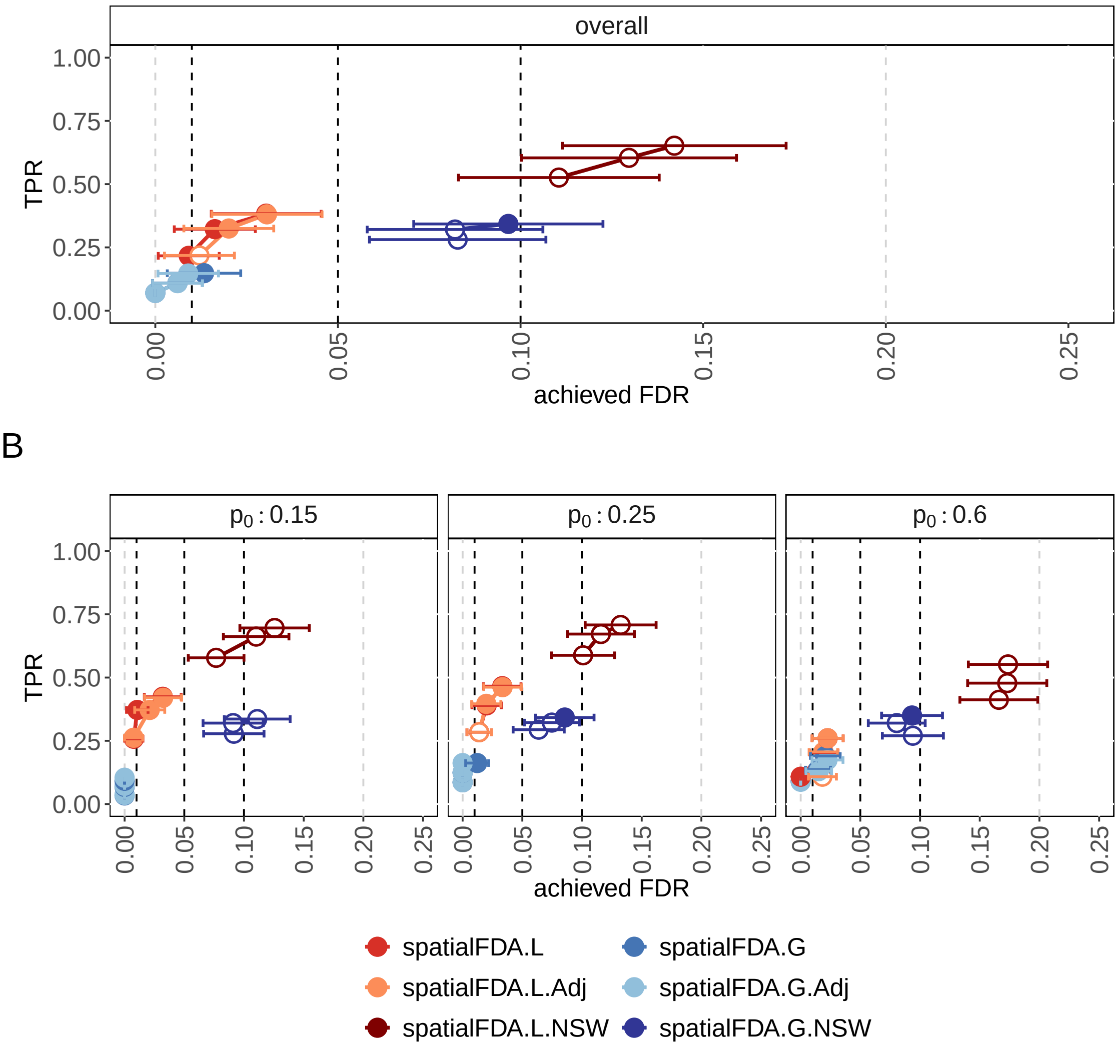


Figure S7: Calibration plots of true-positive rate vs. achieved false discovery rate for all `spatialFDA` variants in the Baker et al. simulation. `spatialFDA.L` and `spatialFDA.G` are not intensity adjusted but have sandwich-corrected covariance matrices. `spatialFDA.L.Adj` and `spatialFDA.G.Adj` have an additional model covariate adjusting for non-spatial intensity differences. `spatialFDA.L.NSW` and `spatialFDA.G.NSW` show the calibration without sandwich corrected covariance matrices. The error bars indicate Wald confidence interval of the Monte Carlo standard errors (MCSE) per observed FDR. **A** Overall performance results across all simulations. **B** Performance results split by the prior cell type proportion of the perturbed cell type.

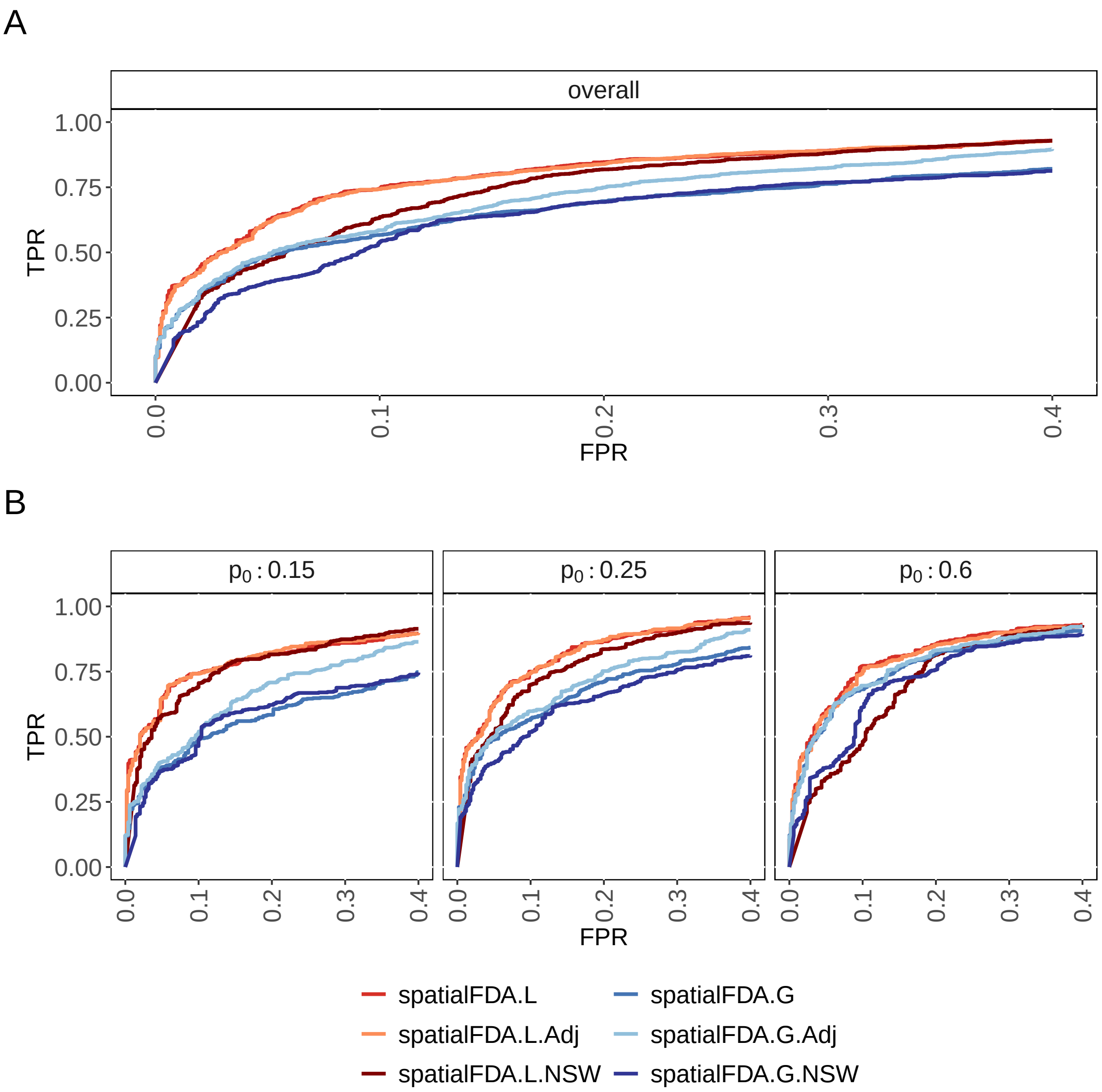


Figure S8: Receiver operating characteristics curve of true-positive rate vs. false-positive rate for all the different `spatialFDA` variants in the Baker et al. simulation. `spatialFDA.L` and `spatialFDA.G` are not intensity adjusted but have sandwich-corrected covariance matrices. `spatialFDA.L.Adj` and `spatialFDA.G.Adj` have an additional model covariate adjusting for non-spatial intensity differences. `spatialFDA.L.NSW` and `spatialFDA.G.NSW` show the performance without sandwich corrected covariance matrices. **A** Overall performance results across all simulations. **B** Performance results split by the prior cell type proportion of the perturbed cell type.

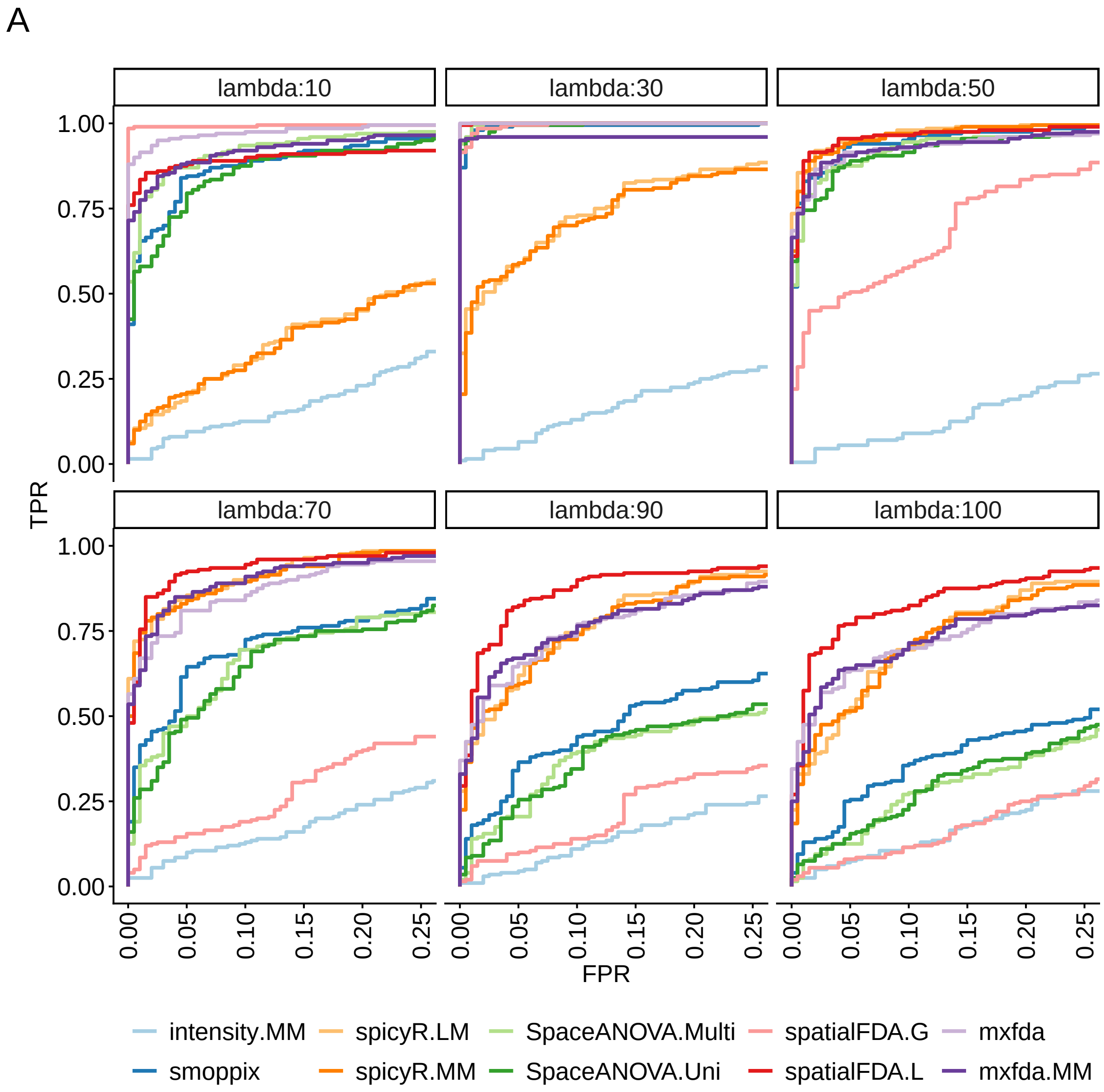


Figure S9: Receiver operating characteristic curves for the simulation from Canete et al. All methods are parameterised to calculate spatial metrics ($L$ and $G$ curves) on a [0,100] interval and compared by ABC (`spicyR`, `smoppix`) or functional data analysis (`spatialFDA`, `mxfda`, `SpaceANOVA`) as well as a non-spatial baseline (`intensity.MM`). The true length scale of the differential CCoL effect changes between the facets of the plot from $10\mu m$ to $100\mu m$ CCoL effect between the two simulated cell types.

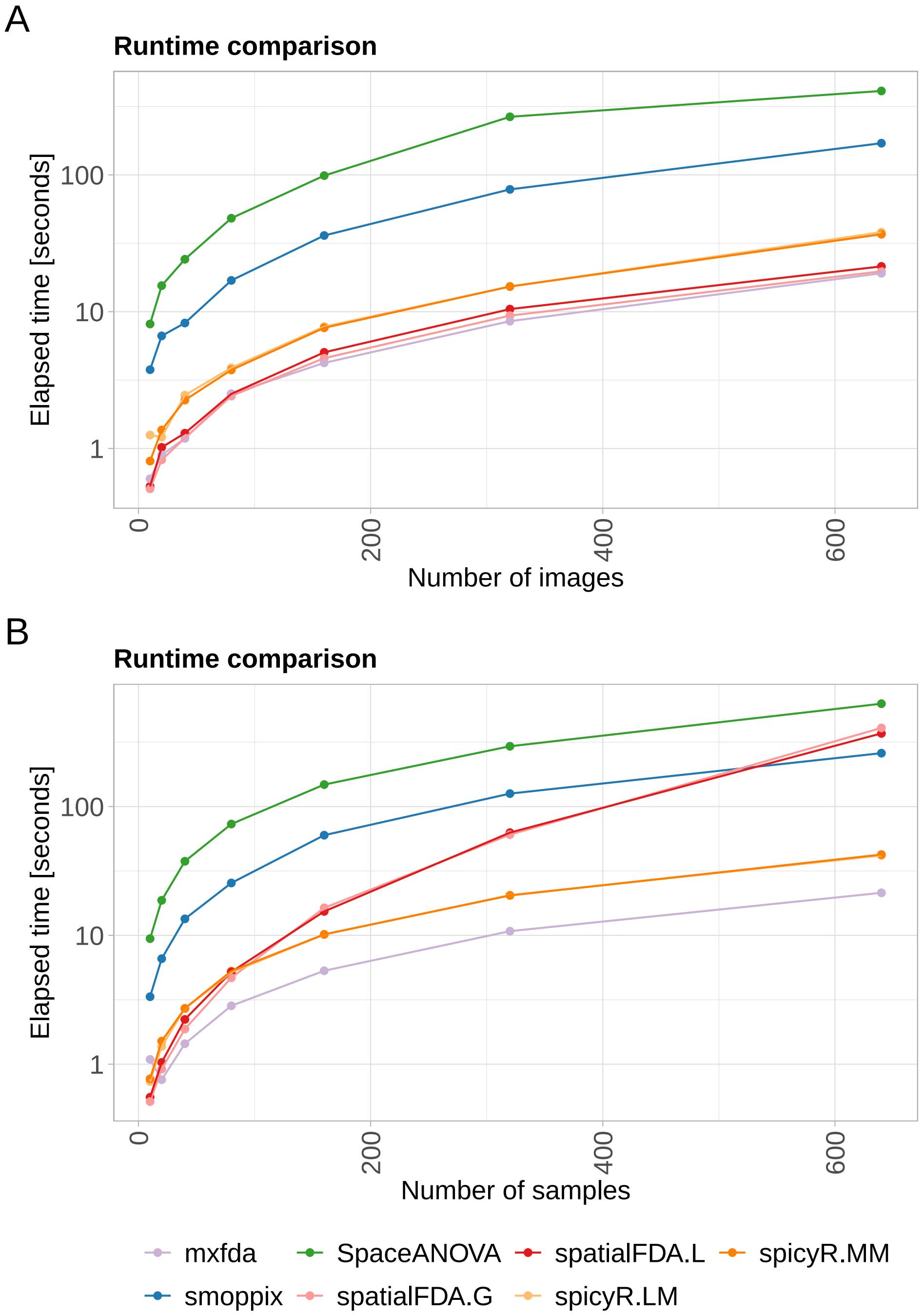


Figure S10: Runtime comparison of all the methods. All methods are run ten times and the average elapsed time over the ten runs is reported. `SpaceANOVA` is only run as one method, this produces both the univariate and multivariate results. `mxfda` was only tested in the fixed-model setting as the mixed model version did not perform well in the TPR/FDP benchmark. **A** Runtime as a function of the number of images. **B** Runtime as a function of the number of independent samples (with multiple images each).

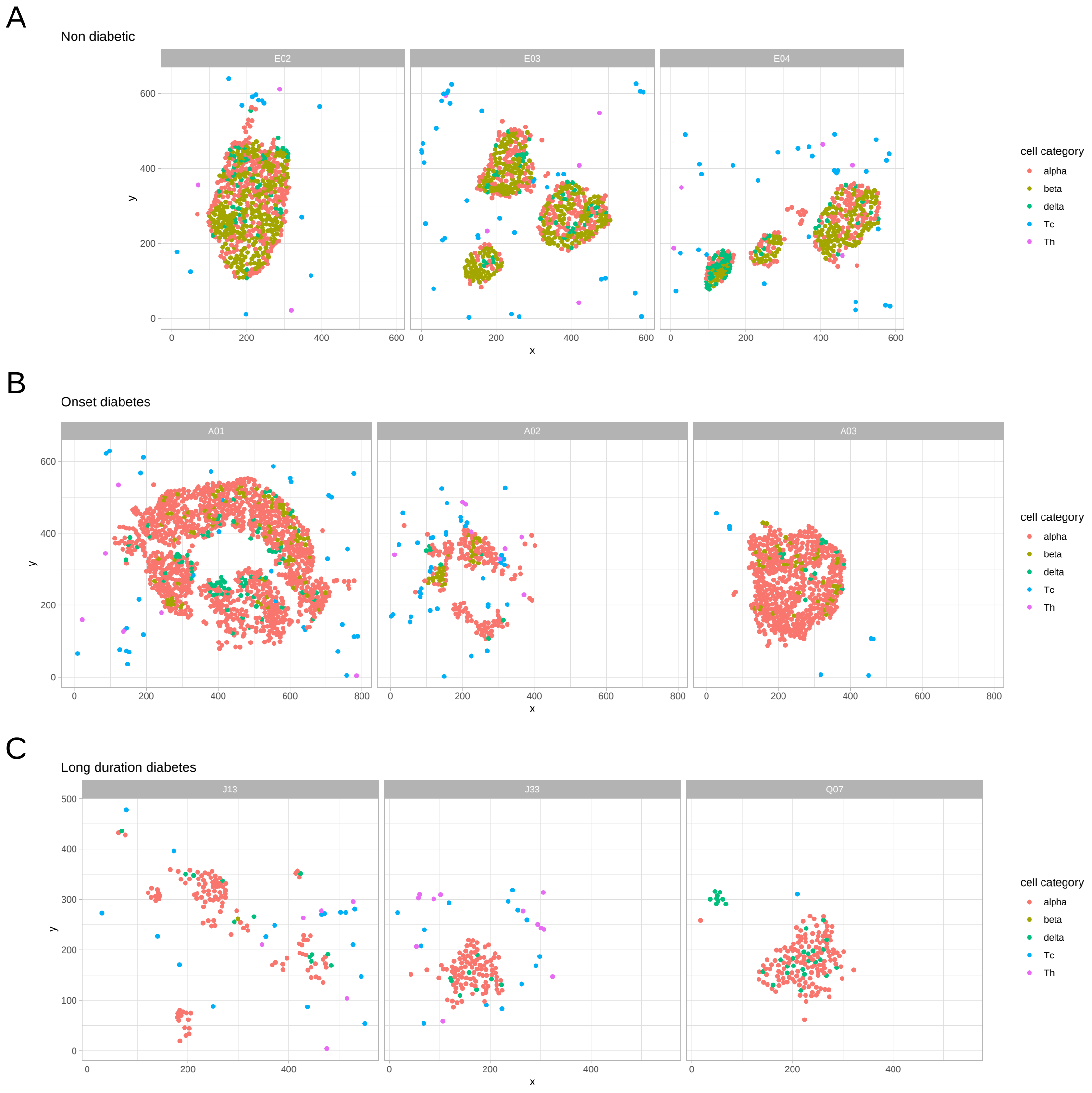


Figure S11: Example fields-of-view for the diabetes case study. Three FOVs are chosen per condition and the cell types are subset to the islet cell types $\alpha, \beta, \delta$ and the two T-cell types, helper (Th) and cytotoxic (Tc). **A** Three non-diabetic fields-of-view, **B** three onset diabetes fields-of-view and **C** three long duration diabetes fields-of-view.

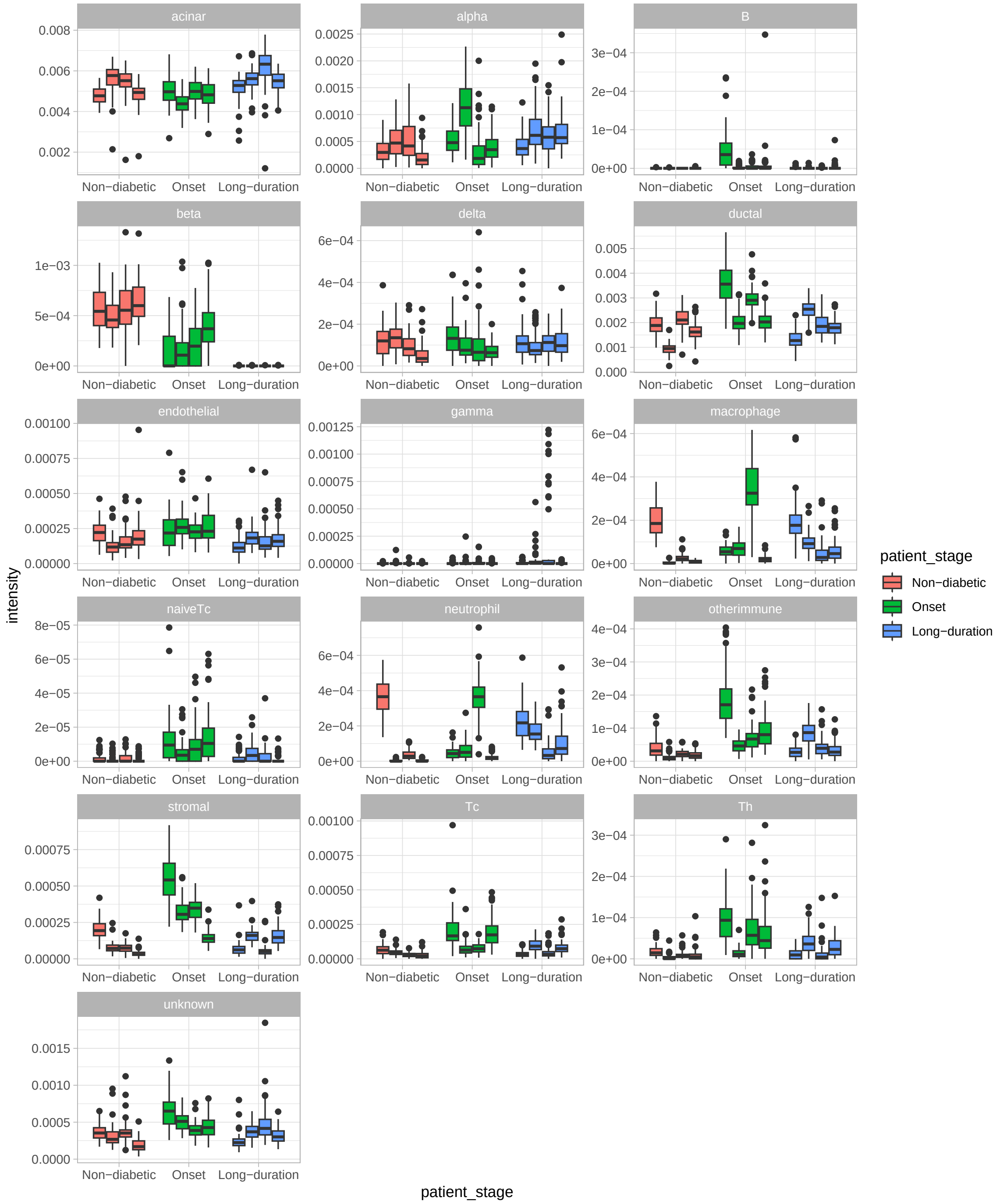


Figure S12: Differences in average intensity per image for the diabetes case study, aggregated by patient as a box plot and repeated for all cell types.

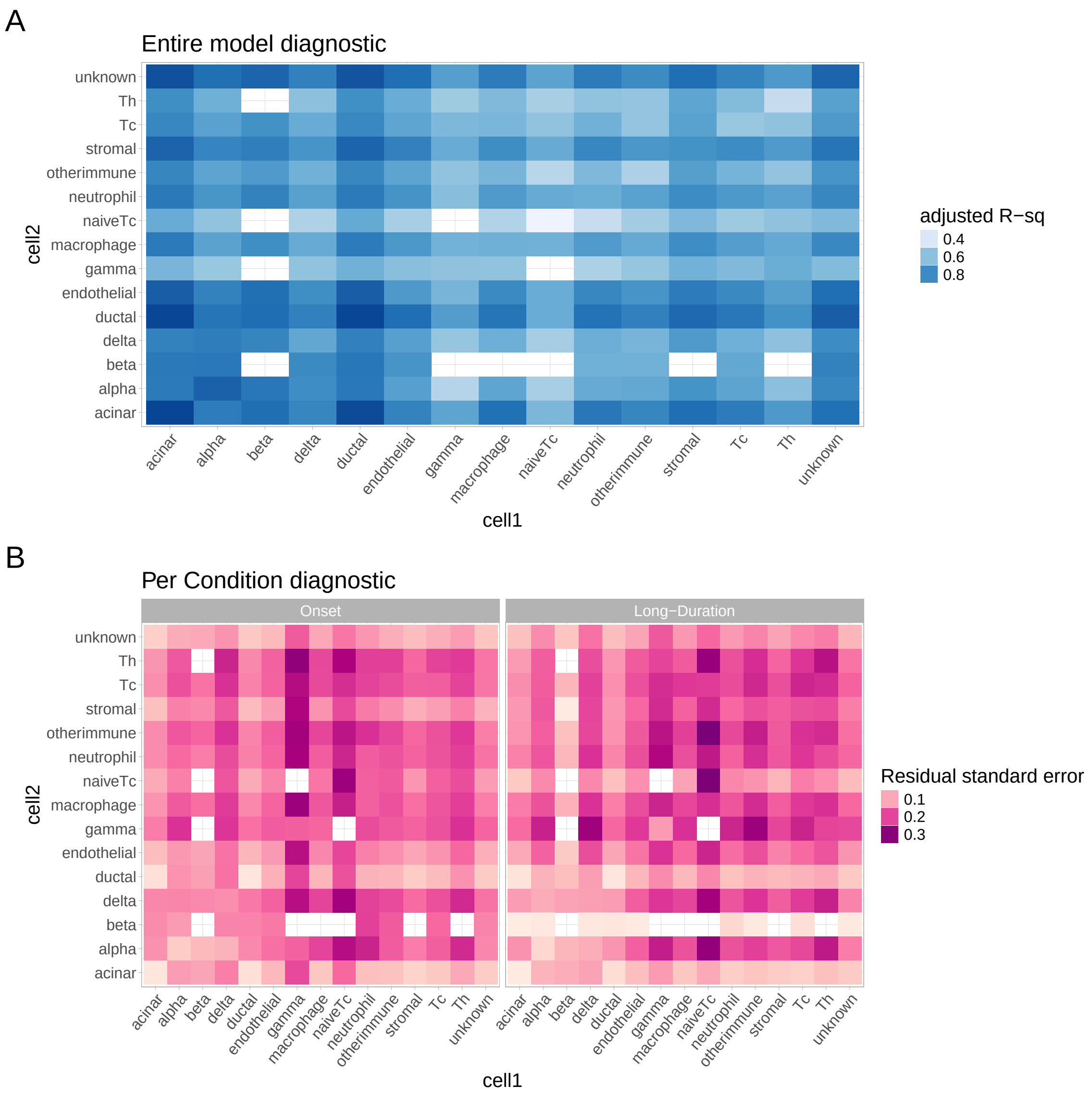


Figure S13: Diagnostic plots of the functional GAM models. **A** is the adjusted R-squared of the entire model fit. The closer this value is to 1, the better. **B** The residual standard error split by condition. The closer this value is to 0, the better.

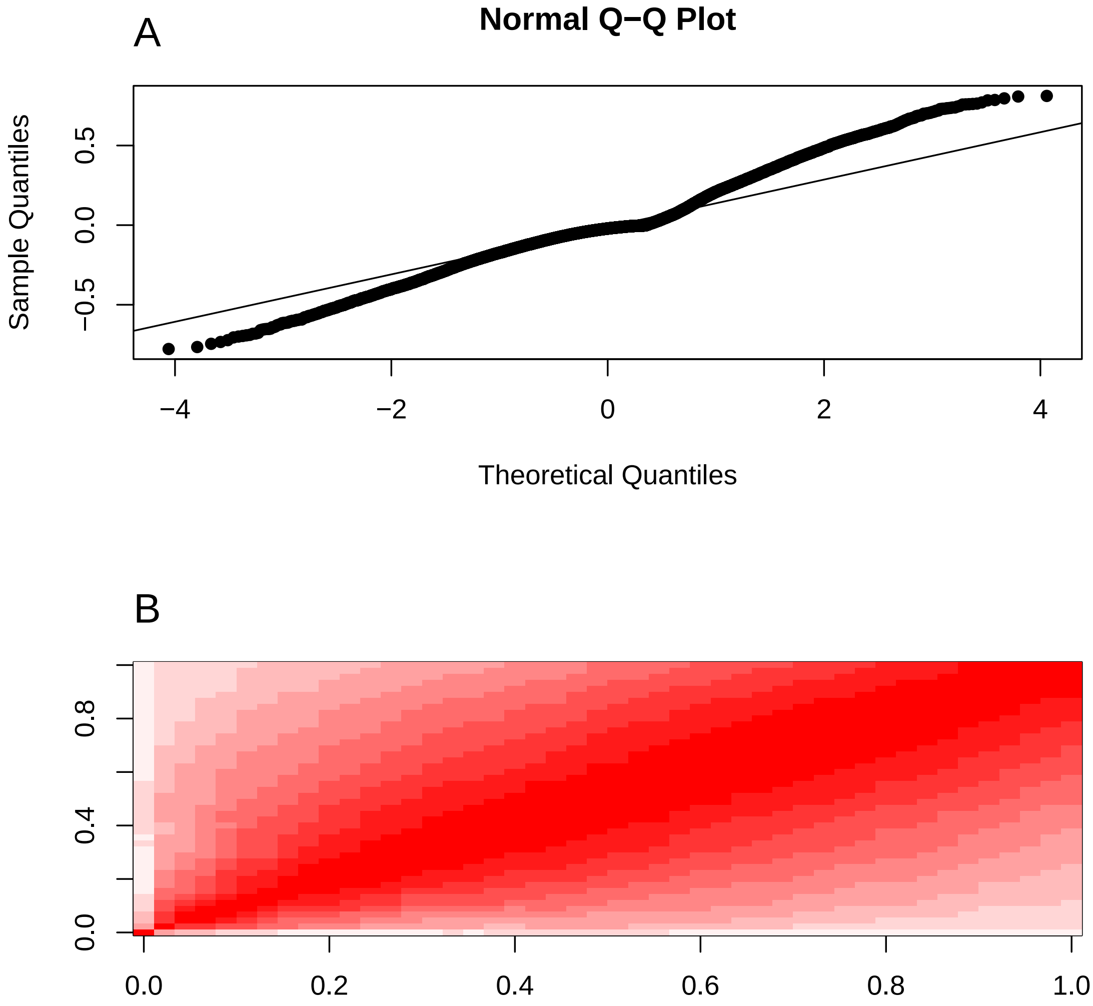


Figure S14: Diagnostic plots of the functional GAM model for the $\delta$-Th cell type interactions. **A** The normal Q-Q plot of the residuals shows whether the residuals of the fGAMM follow normality. **B** The autocorrelation of the residuals along the domain $r$ (scaled to the interval [0,1].

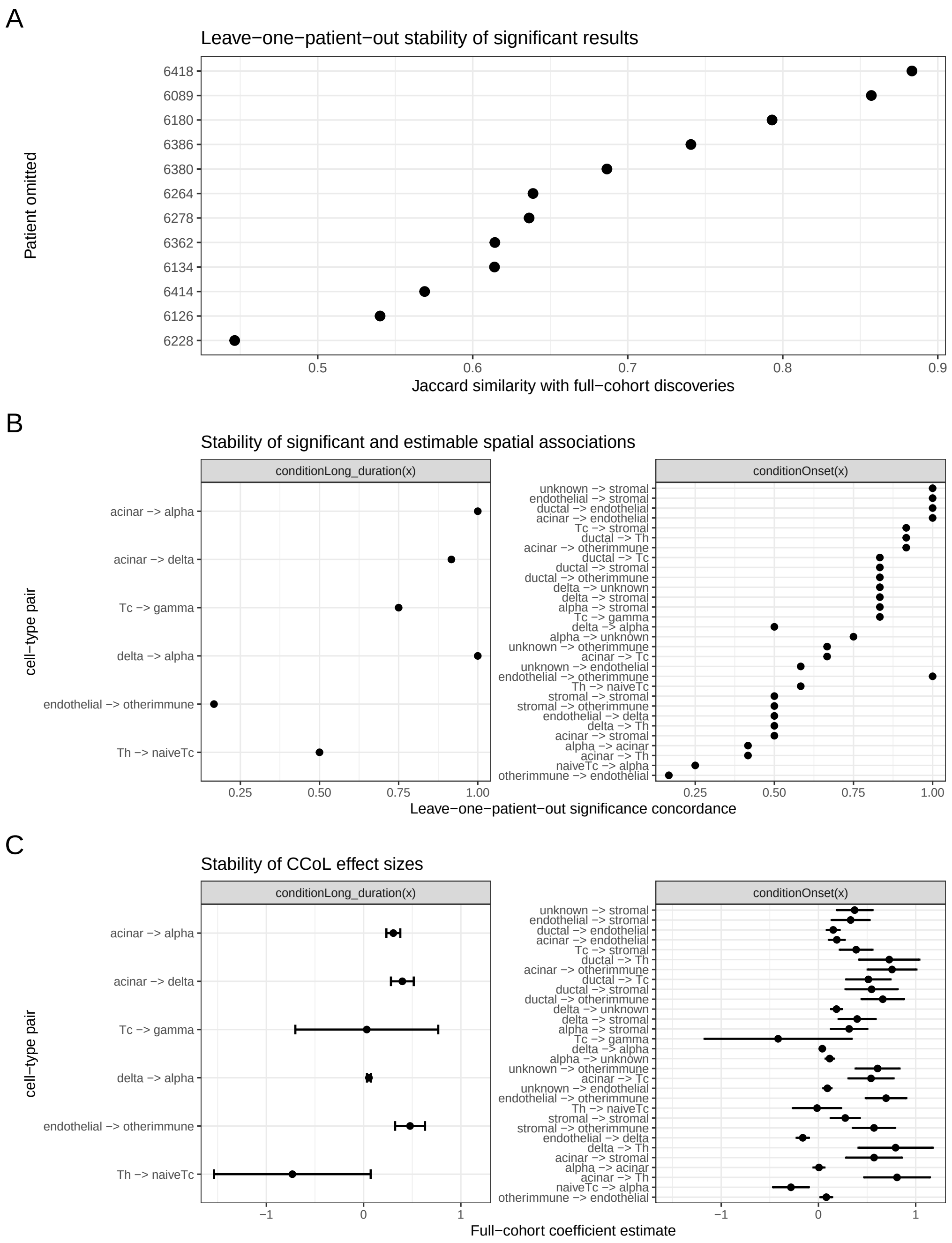


Figure S15: Leave-one-patient-out (LOO) sensitivity analysis of the diabetes analysis results. **A** The Jaccard similarity of the significant results between the full dataset and the dataset without the ommited patient on the $y$-axis. **B** The stability of specific cell type combinations as the concordance rate in which this term remained significant across the LOO folds. This measure is only reported for significant pairs that were estimated in all LOO folds. **C** The stability of the effect size estimate as the mean coefficient over the domain $r$ across LOO folds. The errors represent the standard errors of the mean effect sizes across the LOO folds. This measure is only reported for significant pairs that were estimated in all LOO folds.

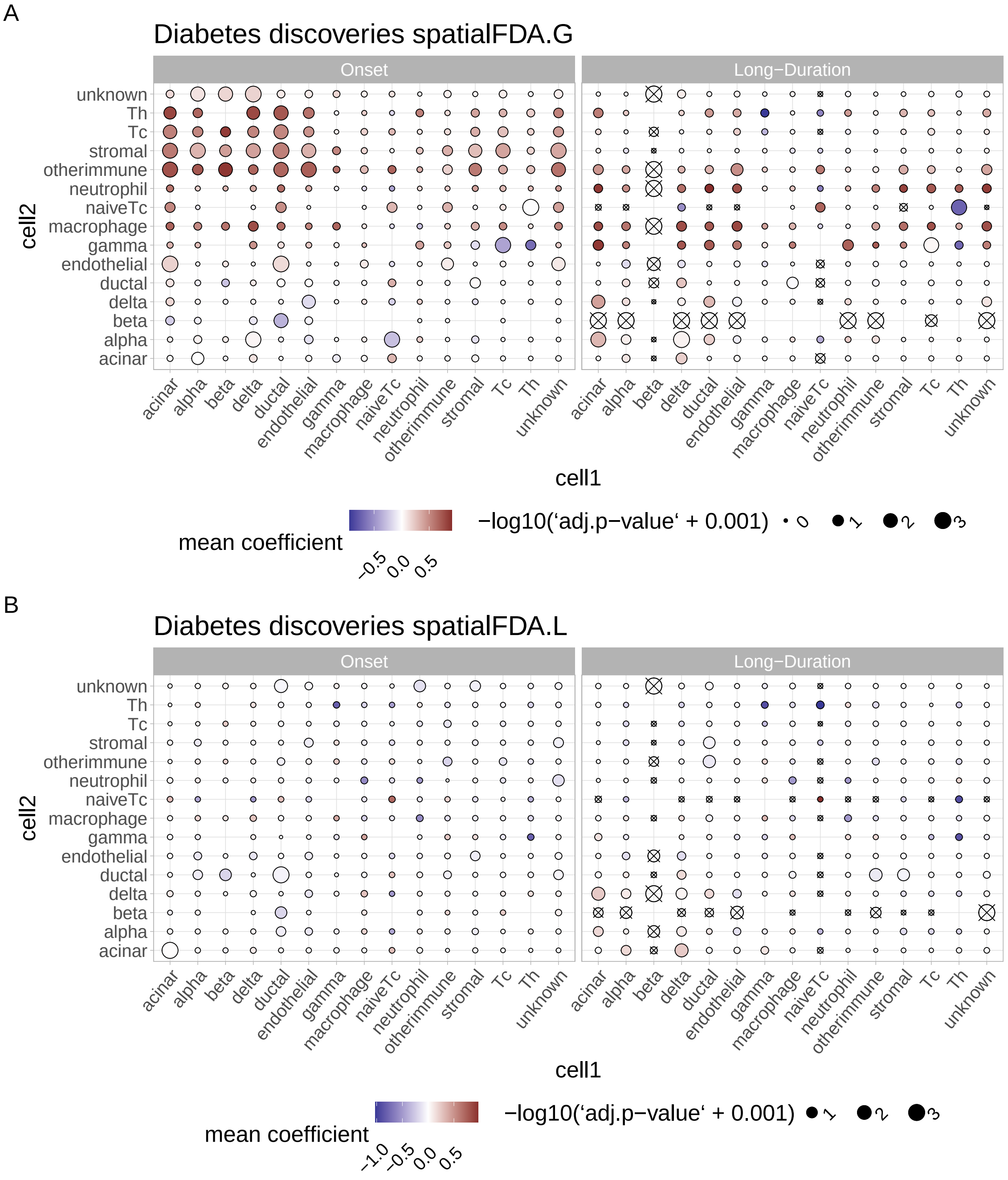


Figure S16: Overall heatmap of the $F$-test results for both `spatialFDA` versions. **A** is with a $G$- and **B** with an $L$-function. The columns are the cell types 1 from which the spatial metric to cell types 2 in the rows is calculated. Colouring indicates the mean functional model value over the functional domain $r$, whereas the size of the circles indicates the negative log $p$-valus corrected for multiple testing (FDR correction [3]).

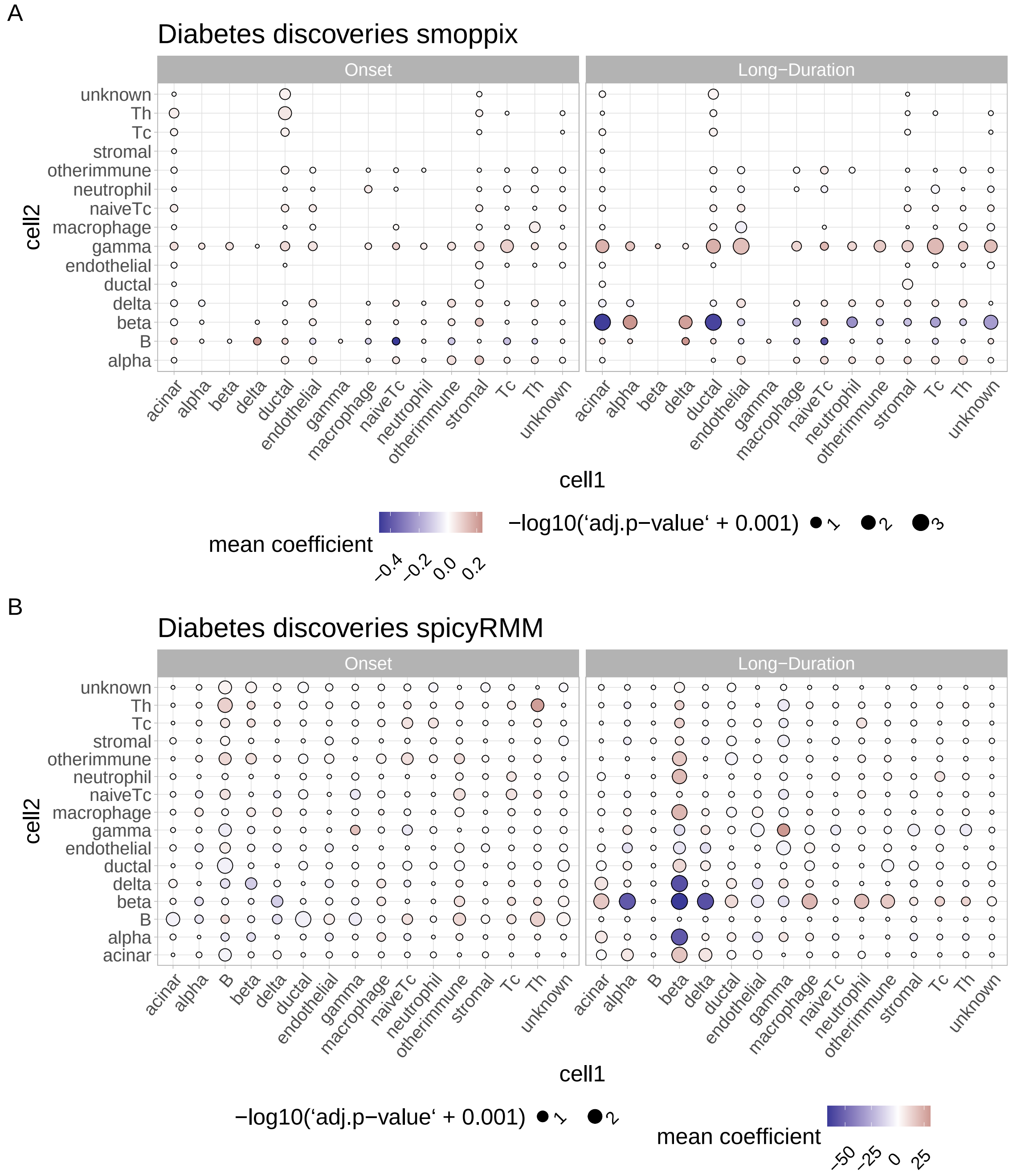


Figure S17: Overall heatmap of the linear mixed effects model results. **A** are the results for `spicyR` and **B** for `smoppix`. The columns are the cell types 1 from which the spatial metric to cell types 2 in the rows is calculated. Colouring indicates the effect size of the coefficient in the linear model whereas the size of the circles indicates the negative log $p$-valus corrected for multiple testing (FDR correction [3]).

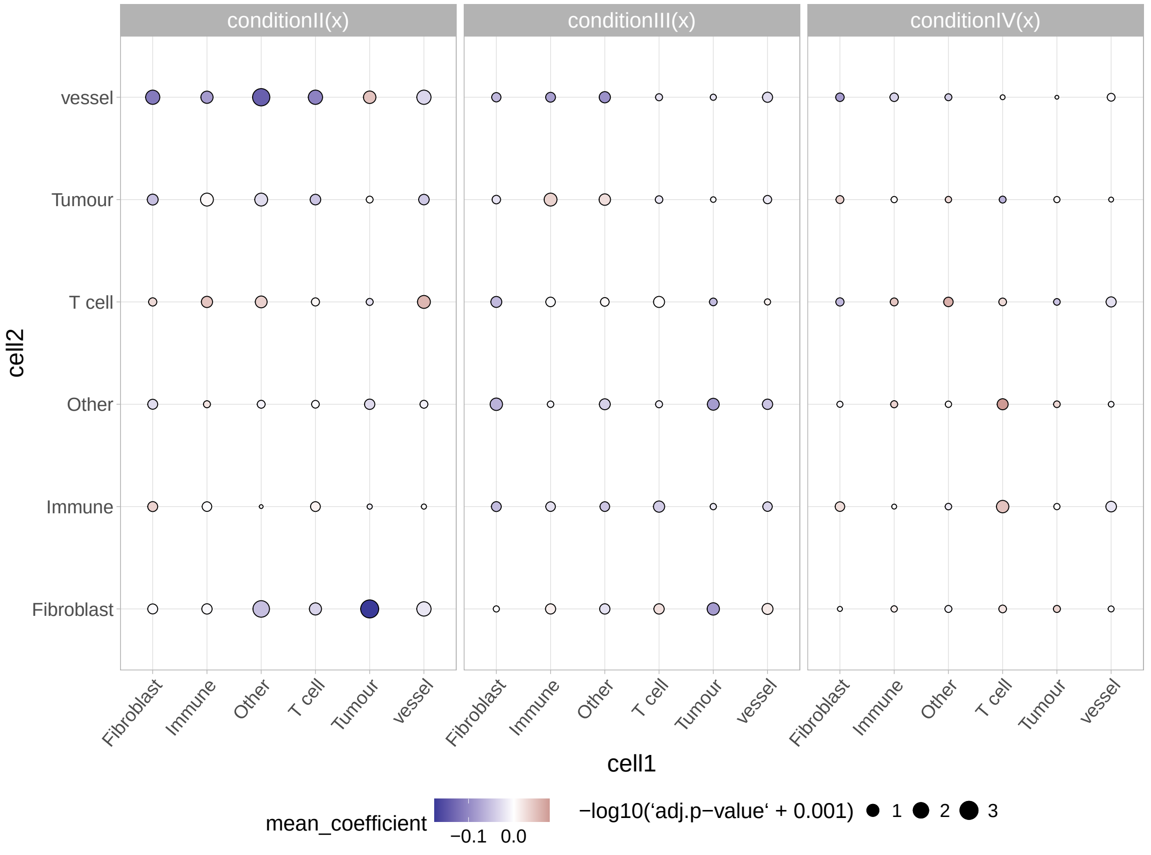


Figure S18: Overall heatmap of differential CCoL results for the Cords et al. study with fixed effects model and choosing one image per patient at random. The colour indicates the mean functional coefficient over the entire domain $r$ whereas the size of the circles is the negative $\log p$-values adjusted for multiple testing (FDR correction [3]).
